# Inflammation induces bitter taste oversensitization via epigenetic changes in *Tas2r* gene clusters

**DOI:** 10.1101/2023.02.08.527520

**Authors:** Cailu Lin, Masafumi Jyotaki, John Quinlan, Shan Feng, Minliang Zhou, Peihua Jiang, Ichiro Matsumoto, Liquan Huang, Yuzo Ninomiya, Robert F. Margolskee, Danielle R. Reed, Hong Wang

**Affiliations:** Monell Chemical Senses Center, 3500 Market St., Philadelphia, PA 19104, USA; Japan Tobacco Inc., Tobacco Science Research Center, 6-2 Umegaoka, Aoba-ku, Yokohama, Kanagawa 227-8512, Japan; University of Maryland School of Medicine and Fischell Department of Bioengineering, University of Maryland, College Park, Maryland, USA; College of Pharmaceutical Sciences and Chinese Medicine, Southwest University, Chongqing, China; Institute of Cellular and Developmental Biology, College of Life Sciences, Zhejiang University, Hangzhou, Zhejiang 310058, China; Division of Sensory Physiology, Research and Development Center for Five-Sense Device, Kyushu University, Fukuoka, Japan; Okayama University, Okayama, Japan; Oral Science Research Center, Tokyo Dental College, Tokyo, Japan

**Keywords:** bitter receptors, chromatin accessibility, epigenetic regulation, taste distortion, chemosensory dysfunction, inflammation, scATAC-seq

## Abstract

T2R bitter receptors, encoded by *Tas2r* genes, are not only critical for bitter taste signal transduction but also important for defense against bacteria and parasites. However, little is known about whether and how *Tas2r* gene expression are regulated. Here we show that, in an inflammation model mimicking bacterial infection, the expression of many *Tas2rs* are significantly up-regulated and mice displayed markedly increased neural and behavioral responses to bitter compounds. Using single-cell assays for transposase-accessible chromatin with sequencing (scATAC-seq), we found that the chromatin accessibility of *Tas2rs* was highly cell type specific and inflammation increased the accessibility of many *Tas2rs*. scATAC-seq also revealed substantial chromatin remodeling in immune response genes in taste tissue stem cells, suggesting potential long-term effects. Together, our results suggest an epigenetic mechanism connecting inflammation, *Tas2r* gene regulation, and altered bitter taste, which may explain heightened bitter taste that can occur with infections and cancer treatments.

## Introduction

T2Rs (encoded by *Tas2rs* in mice or *TAS2Rs* in humans) were identified more than twenty years ago as bitter taste receptors (Adler et al., 2000). However, their functions extend far beyond taste signal transduction. T2Rs expressed in solitary chemosensory cells have been shown to play important roles in recognizing bacterial quorum-sensing molecules and are involved in bacterial defense in the upper airway (Lee et al., 2014; Tizzano et al., 2010; Wang et al., 2021). T2Rs in gut tuft cells are involved in defense against parasites (Luo et al., 2019). Interestingly, thymic tuft cells also express an array of *Tas2r* genes and may be involved in T cell development (Miller et al., 2018). T2Rs have also been implicated in allergies (Grassin-Delyle et al., 2015). These studies suggest that T2Rs are an integral part of immune responses. While it is well known that the genes involved in immunity are often regulated by inflammatory stimuli, whether and how the expression of *Tas2rs* are regulated by inflammation remain to be determined.

Bitter taste is a basic taste quality and believed to serve as a warning signal for potentially harmful compounds in food. To humans and many animal species, bitter taste is innately aversive (Cowart, 2005), even though humans can learn to accept some bitterness for pharmaceutical or nutritional reasons. Bitter taste sensitivity is linked to food preference and intake and influences the consumption of vegetables and alcohol (Duffy et al., 2010; Hayes et al., 2011; Hinrichs et al., 2006). Genetic polymorphisms in *TAS2R* genes contribute to variations in bitter taste sensitivity (Drayna, 2005). Some diseases also appear to affect bitter taste. Heightened or persistent bitter or metallic taste, a type of taste distortions, can occur in patients with infections or undergoing cancer treatments and contributes to food aversion and low quality of life (Schiffman, 1983; Thomas et al., 2022). In traditional Chinese medicine, oral bitterness is an indication used for disease diagnosis. However, the underlying cause for bitter taste oversensitization is unknown.

Embedded in the oral epithelium and exposed to the oral cavity, taste buds are vulnerable to infections (Doyle et al., 2021). Our previous studies showed that taste cells are equipped with proteins in innate immune pathways, including multiple Toll-like receptors (TLRs) for detecting pathogen-derived molecules (Wang et al., 2007; Wang et al., 2009b). Activation of innate immune pathways changes the expression of numerous gene (Medzhitov and Horng, 2009), which is protective against infections but can also lead to pathology. Whether innate immune responses directly regulate *Tas2r* gene expression and bitter taste is unclear.

In this study, we show that lipopolysaccharide (LPS), a pathogen-associated molecular pattern (PAMP) in bacteria and a ligand for TLR4, induces expression of the majority of *Tas2rs* and markedly increases neural and behavioral responses to bitter compounds. Using the single-cell assay for transposase-accessible chromatin with sequencing (scATAC-seq) approach, we show that the genes in *Tas2r* clusters are strongly co-regulated and that LPS increases chromatin accessibility of many *Tas2rs*, indicating an epigenetic mechanism for heightened bitter taste under disease conditions. Our results also reveal that LPS-induced inflammation results in broad changes in the chromatin accessibility landscape in taste tissue stem cells, suggesting potentially long-lasting effects from inflammation.

## Results

### Bitter taste responses were markedly elevated in LPS-induced inflammation

To investigate whether *Tas2r* gene expression and bitter taste are regulated by innate immune responses, we use the well-established systemic inflammation model induced by LPS. Bacterial LPS activates TLR4-mediated signaling pathway and stimulates the production and secretion of an array of inflammatory cytokines (Janeway and Medzhitov, 1999). This inflammatory cascade also occurs in taste tissues (Cohn et al., 2010; Wang *et al*., 2009b). Here, we carried out experiments to measure behavioral and neural responses to taste compounds in this inflammation model (Fig. 1a). As shown in Fig. 1b, we found that LPS-treated mice displayed strong avoidance behavior to the bitter compound quinine compared to mice that received phosphate-buffered saline (PBS; a vehicle control). To confirm that the hyper-avoidance behavior to quinine was due to increased peripheral bitter taste responses, we conducted gustatory nerve recording experiments from both chorda tympani and glossopharyngeal nerves that innervate taste buds in the anterior and posterior parts of the tongue, respectively. Results from these experiments demonstrated that, indeed, LPS treatment induced significantly stronger gustatory nerve responses to bitter compounds (Fig. 1c and Supplementary Fig. 1b). Surprisingly, even though we detected dampened behavioral responses to sucrose in LPS-treated mice in brief-access tests (Supplementary Fig. 1a), the chorda tympani and glossopharyngeal nerve responses to sucrose, as well as to several other sweet compounds, did not show detectable differences in LPS-treated mice compared to control mice (Fig. 1d and Supplementary Fig. 1b). This result suggests that the reduced behavioral responses to sweet compounds were likely due to brain-mediated mechanisms, consistent with previous reports (Dantzer, 2001). We also did not observe significant differences in taste nerve responses to umami or salt compounds (Fig. 1e, f and Supplementary Fig. 1b). LPS-treated mice showed reduced responses to sour compounds only at high concentrations of HCl and citric acid in chorda tympani recordings (Fig. 1f). Together, our results show that LPS-induced inflammation selectively alters bitter taste responses through peripheral mechanisms.

**Fig. 1.**
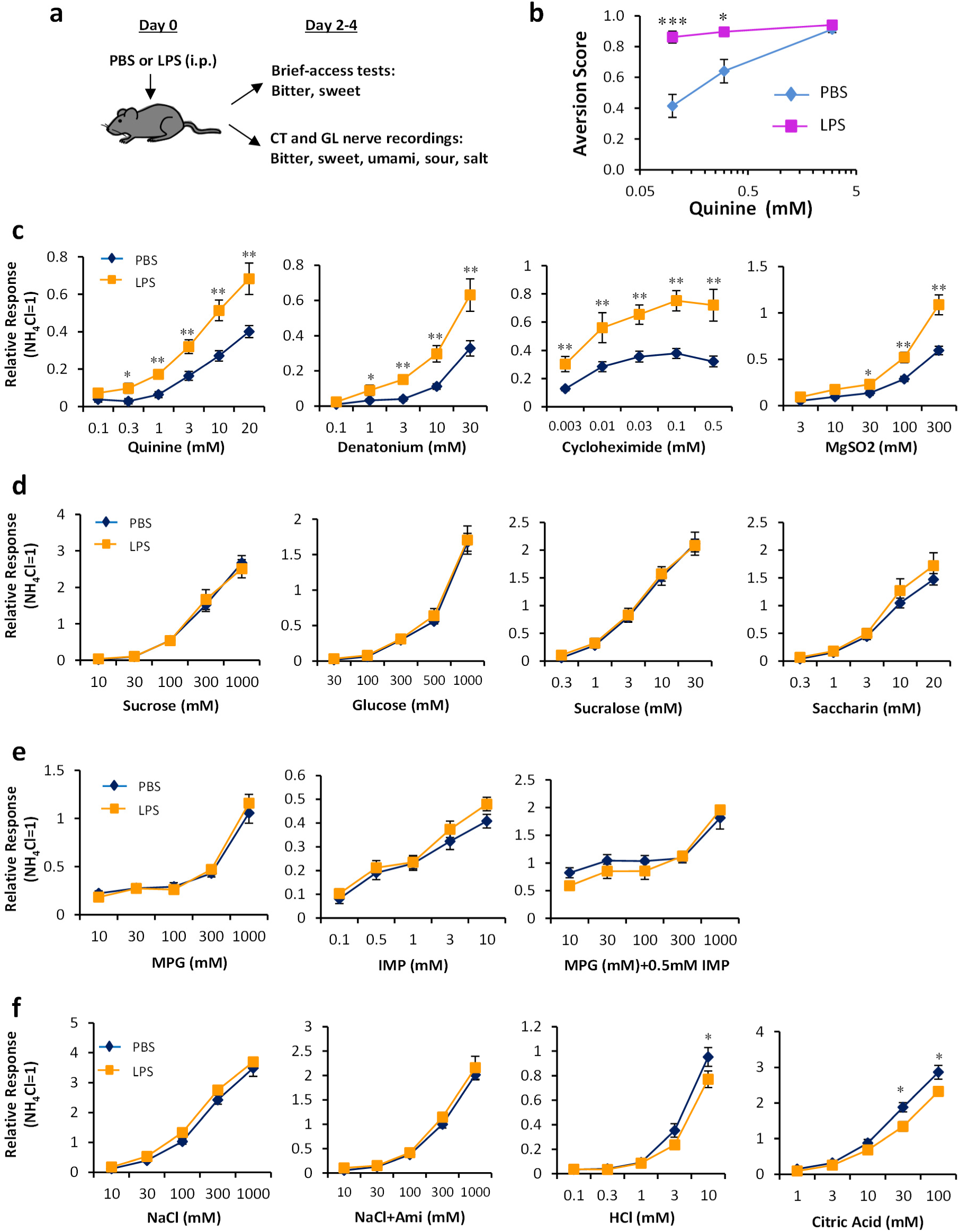
LPS induced neural and behavioral responses to bitter compounds. **a.** Timeline of experiments. CT, chorda tympani nerve; GL, glossopharyngeal nerve. **b.** LPS increased behavioral responses to quinine in brief-access tests. The taste behavioral tests were performed on day 3 after LPS or PBS (control) injection. Aversion scores were calculated as 1 – the lick ratio (tastant licks/water licks). * *p*<0.05; *** *p*<0.005 (ANOVA with post hoc t tests). N=8-10 mice per group. **c-f.** LPS selectively increased chorda tympani nerve responses to bitter compounds (**c**) but not to sweet (**d**), umami (**e**), or salty or sour compounds (**f**). MPG, monopotassium glutamate; IMP, inosine-5’-monophosphate; Ami, amiloride. Gustatory nerve recordings were conducted on day 3 after PBS or LPS injection. Data are means ± SE. * *p*<0.05; ** *p*<0.01 (ANOVA with post hoc t tests). N=6-12 mice per group.

### LPS stimulated the expression of the majority of *Tas2rs*

To investigate the underlying mechanism for the LPS-induced hyper-responsiveness to bitter compounds, we analyzed *Tas2r* gene expression in the taste epithelium from circumvallate and foliate taste papillae by real-time quantitative reverse-transcription polymerase chain reaction (qRT-PCR). As shown in Fig. 2a, LPS treatment stimulated expression of the majority of *Tas2rs*. On day 3 after LPS injection, expression of more than 20 mouse *Tas2rs* was significantly up-regulated by at least 2-fold. We selected several *Tas2rs*, including *Tas2r102*, *108*, *110*, *116*, *124*, and *137*, and analyzed their expression across several time points after LPS injection. Elevated expression of *Tas2R102*, *110*, *116*, and *137* peaked on day 3 after LPS injection (Fig. 2b), whereas expression of *Tas2r108* and *124*, as well as *Trpm5*, peaked on day 5 (Fig. 2c). Expression of *Nfkb1*, a transcription factor involved in inflammatory responses, was also mildly elevated on days 1-5 and returned to control level on day 10 (Fig 2c). Interestingly, 8 of the 10 highly induced *Tas2rs* (>3-fold induction) are located in a closely linked region on mouse chromosome 6 (Fig. 2d), suggesting the transcription of these *Tas2rs* may be co-regulated.

**Fig. 2.**
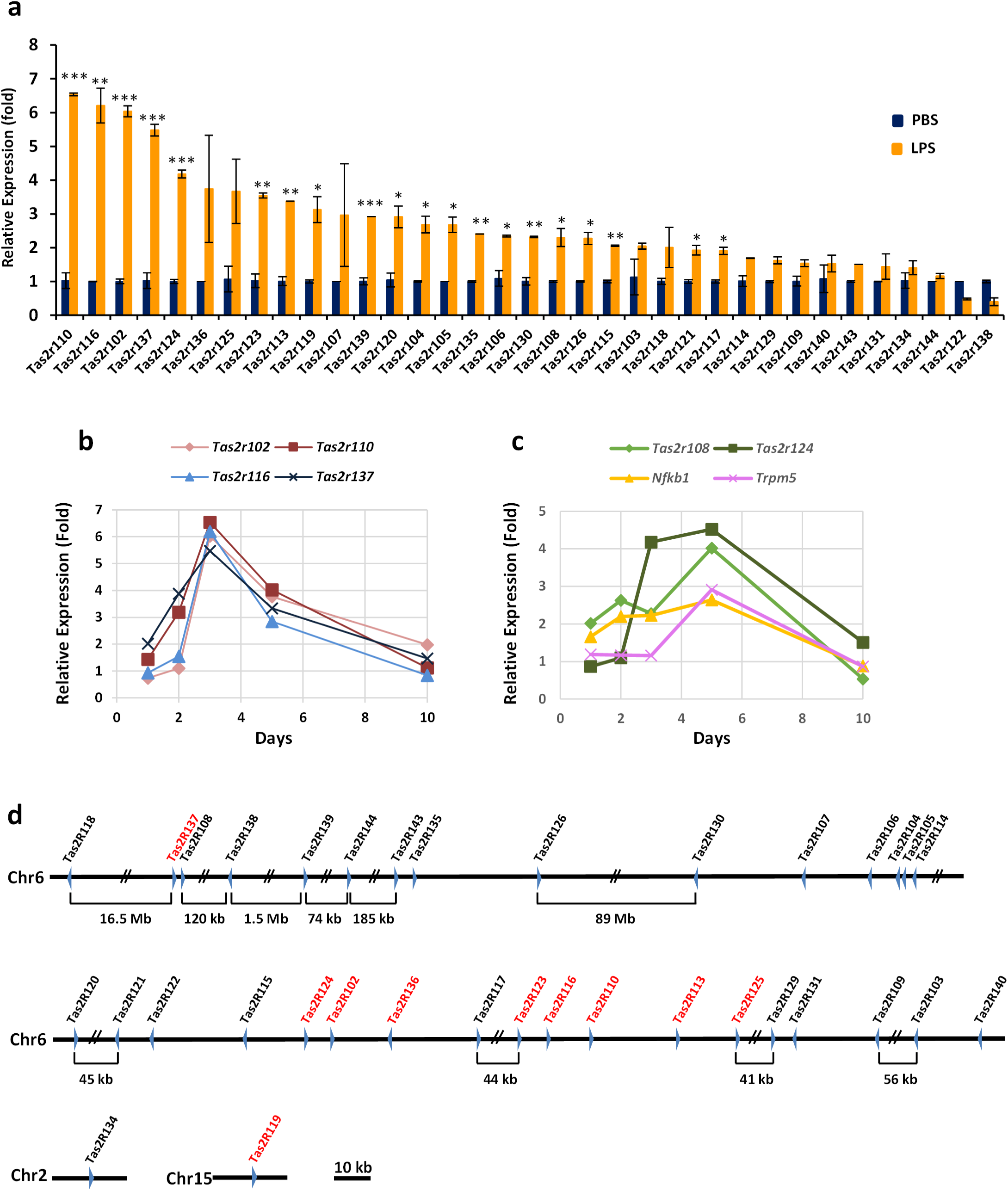
LPS stimulated expression of the majority of *Tas2rs*. **a.** qRT-PCR analysis of mRNA levels of 35 mouse *Tas2rs* in the circumvallate and foliate epithelium 3 days after LPS or PBS injection. Gene expression in PBS samples was set to 1. β-Actin was used as the endogenous control. Data are means ± SE. * *p*<0.05; ** *p*<0.01; *** *p*<0.005 (t tests). **b, c.** Time course of LPS-induced expression of six *Tas2rs*, *Nfkb1*, and *Trpm5*. qRT-PCR experiments were performed as described in **a**. **d.** Schematic illustration of *Tas2r* distribution on mouse chromosomes. Red indicates that the gene was induced >3-fold by LPS. Blue arrowheads indicate direction of *Tas2r* gene transcription.

### scATAC-seq identified taste and nontaste cell clusters and revealed unexpected immune regulatory signature in von Ebner gland duct cells

scATAC-seq can reveal gene regulatory mechanisms, but its use for studying taste receptor gene expression has not been reported. To better understand *Tas2r* gene regulation, we performed scATAC-seq experiments. We isolated single-cell nuclei from the circumvallate epithelium of PBS- (vehicle control) and LPS-treated mice. The scATAC-seq data sets were processed using the ArchR package, which provided comprehensive features for analyzing scATAC-seq data (Granja et al., 2021). Quality checks of the data are shown in Supplementary Fig. 2. The overall quality and the Simpson diversity index of the data sets from the control (PBS) and LPS-treated mice are comparable. In total, 3424 and 4032 nuclei from PBS- and LPS-treated mice, respectively, were included in downstream analyses (Supplementary Fig. 2e). With the combined data sets, we performed uniform manifold approximation and projection (UMAP) dimension reduction and clustering and identified 11 cell clusters (Fig. 3a). The number and percentage of cells in each cell cluster are listed in Supplementary Table 1.

**Fig. 3.**
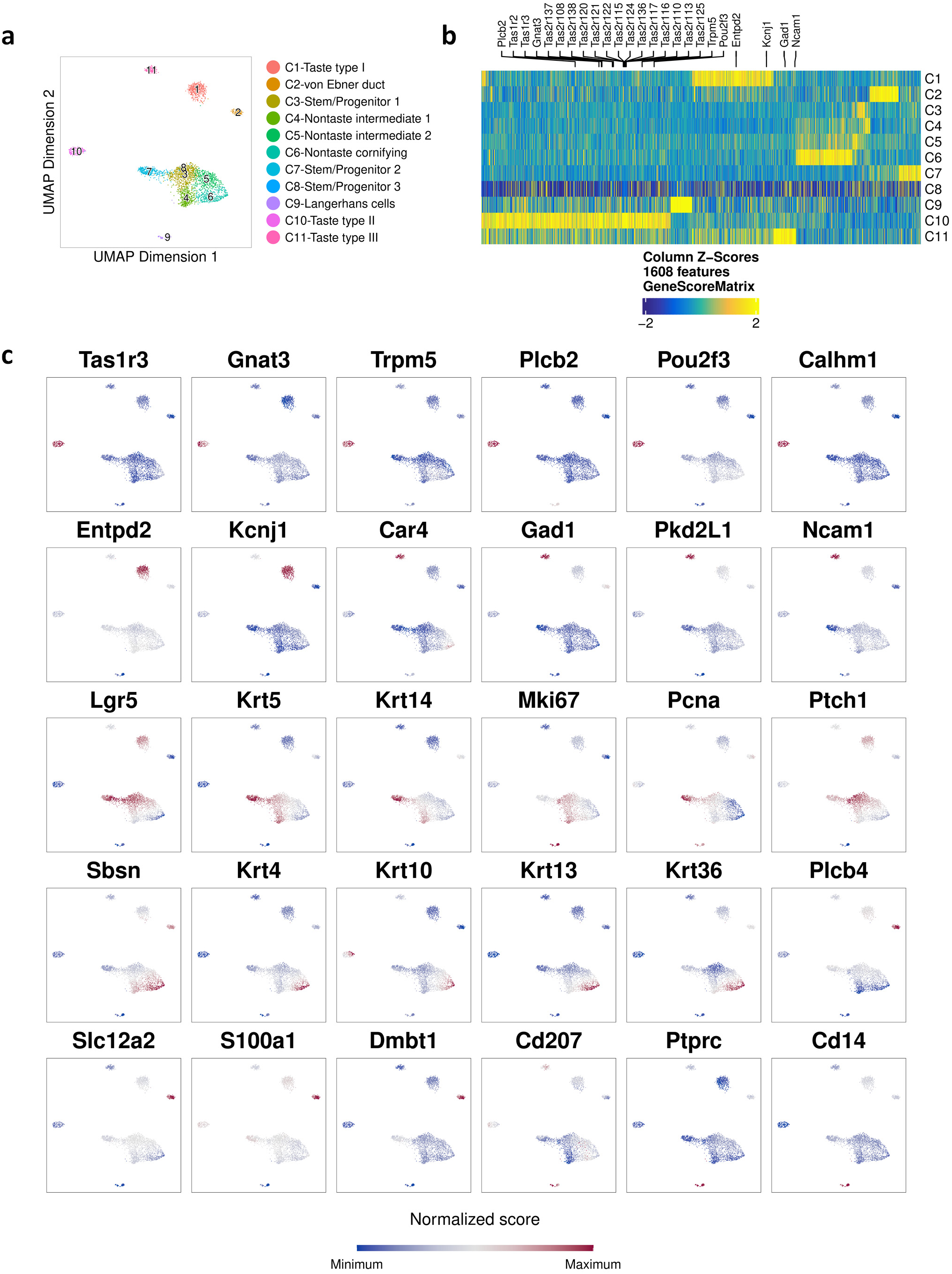
scATAC-seq identified taste and nontaste cell clusters. **a.** UMAP plot of cell clusters of circumvallate epithelia from control mice. The inferred identities of the cell clusters are shown on the right. **b.** A heatmap of gene scores of identified significant marker genes across cell clusters. Some well-characterized marker genes for type I, II, and III taste bud cells are shown on top. **c.** Feature plots of cell-type marker genes that show gene scores in cell clusters. Gene names are shown on the top of each plot.

Next, we performed in-depth analysis of the data set from control mice. We calculated gene scores (or gene activity scores) based on chromatin accessibility using algorithms in ArchR (Granja *et al*., 2021). Chromatin accessibility of individual genes strongly correlates with, and thus can be used as a proxy for, gene expression (Granja *et al*., 2021; Ziffra et al., 2021). To infer the identities of the cell clusters, we generated a gene score heatmap (Fig. 3b) and identified marker genes for each cell cluster. The UMAP plots for well-characterized cell-type markers are shown in Fig. 3c.

Three well-separated cell clusters on the UMAP (Fig. 3a), C1, C10, and C11, showed high gene scores for many taste bud cell markers corresponding to taste cell types I, II, and III, respectively (Finger, 2005; Roper and Chaudhari, 2017). *Entpd2* and *Kcnj1* encode type I taste cell markers (Bartel et al., 2006; Dvoryanchikov et al., 2009) and showed the highest gene scores in the C1 cluster (Fig. 3b, c and Supplementary Data 1), indicating this cell cluster represents type I taste cells. *Gad2* is also enriched in the C1 cluster (Supplementary Data 1), consistent with a recent report showing selective expression of this gene in type I taste cells (Rodriguez et al., 2021). Similarly, many type II taste cell marker genes, such as *Tas1rs*, *Tas2rs*, *Gnat3*, *Trpm5*, *Plcb2*, *Pou2f3*, and *Calhm1* (Roper and Chaudhari, 2017), had the highest gene scores in the C10 cluster (Fig. 3b, c), indicating the C10 cluster denotes type II taste receptor cells. *Car4*, *Pkd2l1*, *Gad1*, and *Ncam1* encode well-established type III taste bud cell markers (Chandrashekar et al., 2009; Finger, 2005), and all showed highest gene scores in the C11 cluster (Fig. 3b, c). Among the three clusters, the percentages of cells in C1 (type I), C10 (type II), and C11 (type III) were 61.6%, 23%, and 15.4%, respectively. This relative abundance of cells was similar to previously reported proportions of type I, II, and III cells in taste buds (Matsumoto et al., 2011; Ogata and Ohtubo, 2020), further supporting our annotations of these clusters. Moreover, this analysis identified many candidate genes that showed selective gene scores in the three types of taste bud cells (Supplementary Data 1).

The C2 cluster is also a well-separated cell cluster on UMAP and shows high gene scores for marker genes of salivary glands. The von Ebner gland, a minor salivary gland, secretes saliva that drains into the circumvallate trenches through gland ducts. In preparing circumvallate epithelium for scATAC-seq, the top portions of the von Ebner gland ducts were also included. Our analysis showed that the C2 cluster was enriched with *Slc12a2*, *S100a1*, *Plcb4*, and *Dmbt1* (i.e., Ebnerin) (Fig. 3c); all have been shown to be selectively expressed in salivary glands (Hauser et al., 2020; Li and Snyder, 1995). *Dmbt1* was shown to be abundantly expressed in von Ebner gland duct cells (Asano-Miyoshi et al., 1998; Li and Snyder, 1995), indicating the C2 cluster represents von Ebner gland duct cells.

The small C9 cell cluster showed high gene scores for CD207 (i.e., Langerin), a well-established marker for Langerhans cells (i.e., epidermis-resident macrophages) (Doebel et al., 2017). *Ptprc* (encodes CD45) and *Cd14* are marker genes for hematopoietic-derived cells and macrophages, respectively, and both showed high gene scores in C9, consistent with the identity of Langerhans cells (Fig. 3c).

Collectively, *Lgr5*, *Krt5*, *Krt14*, and *Ptch1*, genes that encode markers for taste stem/progenitor cells, showed high gene scores in C3, C7, and C8 clusters (Fig. 3c), indicating these clusters represent stem/progenitor cells in the taste tissue. Both *Mki67* and *Pcna* are cell proliferation marker genes, but *Mki67* is more specific for actively proliferating cells (Bologna-Molina et al., 2013; Muskhelishvili et al., 2003). Considering the relative gene scores of the above markers, C7 most likely represents a stem cell cluster due to its high gene score for *Lgr5* but medium score for *Mki67*. C3 and C8 most likely represent transient amplifying cell clusters with high gene scores for *Mki67* and *Ptch1* but medium scores for *Lgr5* (Fig. 3c). Only a small number of cells in control mice were assigned to the C8 cluster, but LPS treatment dramatically increased the number of cells in this cluster (Supplementary Table 1). These data suggest that the C8 cluster could represent a transient cell state between taste progenitor cells and postmitotic taste precursor cells. During normal taste bud turnover, the transition between the cell states proceeds swiftly. However, this transition may be suppressed by LPS-induced inflammation, consistent with our previous reports that showed LPS treatment inhibited taste bud cell renewal (Cohn *et al*., 2010). Furthermore, we found that *Ugt2a1*, a candidate gene related to COVID-19-induced taste and smell loss (Shelton et al., 2022), is enriched in the C3 cluster of stem/progenitor cells (Supplementary Data 1), suggesting a possible role of *Ugt2a1* in taste cell turnover and renewal.

Clusters C4, C5, and C6 showed increasing enrichment for markers of cornifying, terminally differentiating keratinocytes (Supplementary Data 1). The UMAP plots for some of the marker genes, including *Sbsn*, *Krt4*, *Krt10*, *Krt13*, and *Krt36* (Eckhart et al., 2013; Fuchs and Raghavan, 2002; Miura et al., 2014), are shown in Fig. 3c. Cells in the C4 cluster also exhibited relatively high gene scores for *Krt5*, *Krt14*, and *Mki67*, indicating these cells may still have proliferating capacity. Conversely, the C6 cluster displayed high gene scores for the above-mentioned terminal differentiation markers of keratinocytes and most likely represents cornifying nontaste epithelial cells, while the C5 cluster showed an intermediate phenotype (Fig. 3c).

With the identified significant marker genes for each cell cluster, we performed pathway enrichment analyses using g:Profiler (Reimand et al., 2016) and Cytoscape (Shannon et al., 2003)/EnrichmentMap (Merico et al., 2010) software. As expected, type II taste cells showed strong pathway enrichment for GPCR-mediated taste perception, while type III cells were enriched with synaptic activities (Supplementary Fig. 3b, c), consistent with the functions of these taste cells. Interestingly, type I taste cells displayed enrichment for protein secretion, hormone secretion, and hormone transport (Supplementary Fig. 3a), suggesting these cells may play important physiological roles in taste buds, such as connecting hormonal responses to taste sensation.

Surprisingly, von Ebner gland duct cells showed strong enrichment of immune regulatory pathways (Supplementary Fig. 3d). In particular, this cluster was enriched with marker genes encoding chemokines (e.g., *Ccl3*, *Ccl5*, *Ccl6*, *Ccl9*, and *Cx3cl1*) that play critical roles in the migration of various types of immune cells, including neutrophils, monocytes, and lymphocytes (Hughes and Nibbs, 2018). This cluster was also enriched with cytokine response genes, such as those involved in responses to tumor necrosis factor (TNF) and interleukin (IL)-1. Dclk1, a marker for tuft cells, is also a significant marker gene for this cell cluster (Supplementary Data 1). These results suggest that von Ebner gland duct cells may play important roles in immune surveillance of the posterior part of the tongue.

### scATAC-seq revealed cell-type-specific chromatin accessibility of *Tas2r* genes

Chromatin accessibility represents a fundamental feature of cell identity. To date, the chromatin accessibility landscape of taste bud cells has not been reported. To gain insights into cell-type-specific gene expression regulation, we performed chromatin-accessible peak calling using ArchR. Figure 4a shows peak distribution in different genomic regions. Most peaks are located in the intronic and distal regions, followed by the promoter regions, while the exonic regions have the least number of peaks. This pattern of peak distribution is very similar to that observed in other types of tissues, such as retina and brain (Finkbeiner et al., 2022; Ziffra *et al*., 2021).

**Fig. 4.**
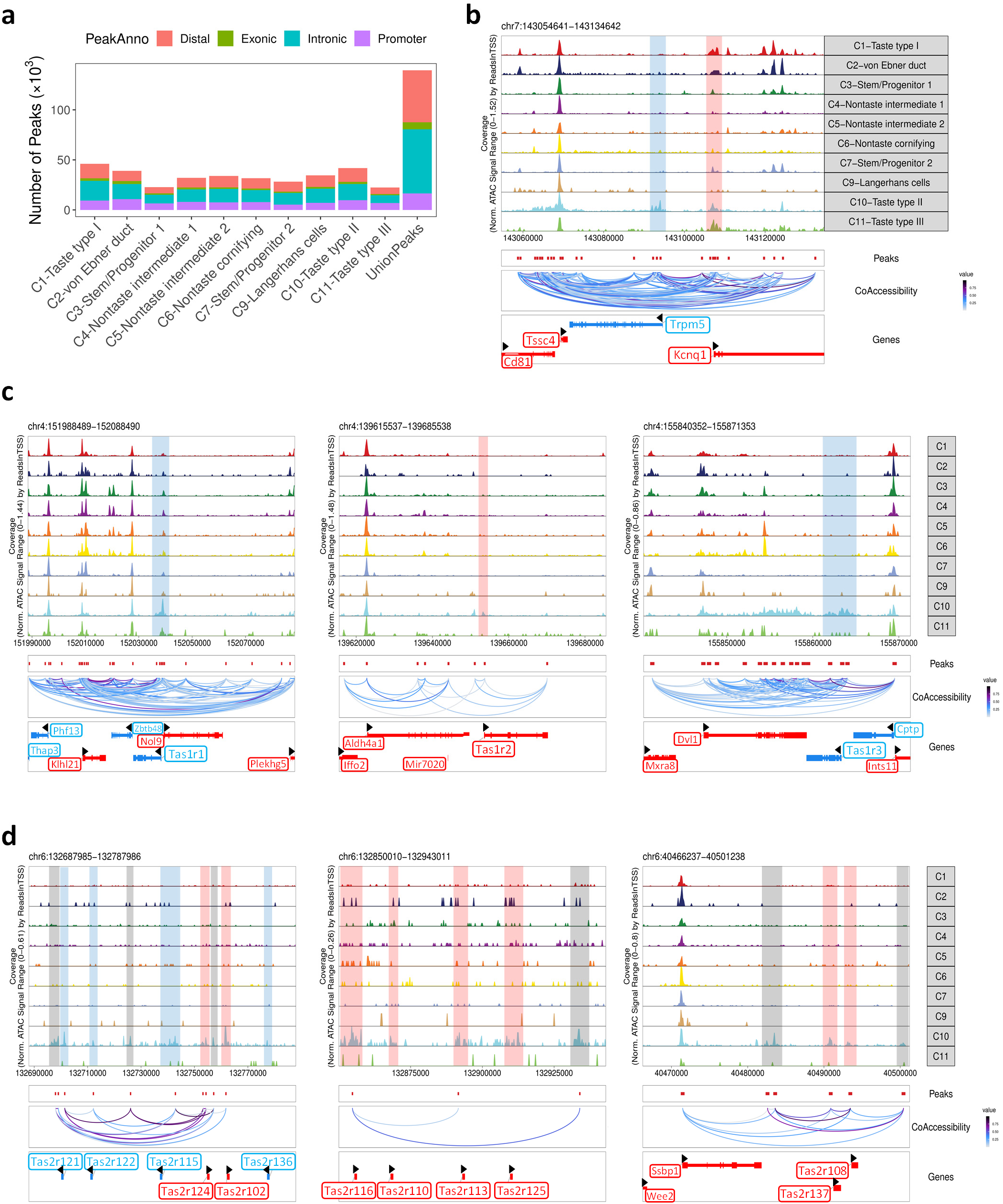
scATAC-seq identified putative regulatory regions in taste receptor and signaling genes. Analysis of chromatin-accessible peaks from control mice. **a.** Distribution of accessible peaks in distal, exonic, intronic, and promoter regions. PeakAnno, peak annotation. **b-d.** Chromatin-accessible peaks in the genomic regions of *Trpm5* and *Kcnq1* (**b**), *Tas1rs* (**c**), and some *Tas2rs* (**d**). Gene symbols in red indicate genes transcribed in the forward direction, and their cluster-selective accessible peaks are highlighted in pink. Gene symbols in blue indicate genes transcribed in the reverse direction, and their cluster-selective accessible peaks are highlighted in blue. Gray bars in **d** highlight type II cell-selective peaks in intergenic regions. Black arrowheads also indicate direction gene transcription. TSS, transcription start site.

The *Tas2r* bitter receptor genes form several gene clusters in the genome (Fig. 2d), and the chromatin accessibility landscapes of some *Tas2rs* are shown in Fig. 4d and Supplementary Fig. 4a. *Tas2rs* showed highly selective accessibility in type II taste cells. Moreover, we identified several intergenic peaks in the *Tas2r* genomic regions that showed selective accessibility in type II cells (gray bars, Fig. 4d and Supplementary Fig. 4a). These peaks and the neighboring *Tas2rs* peaks share strong co-accessibility, suggesting these intergenic peaks could contain putative cis-regulatory elements, such as enhancers for *Tas2rs*.

We also analyzed chromatin accessibility of other well-known genes involved in taste reception and signaling. Figure 4b shows chromatin-accessible peaks for *Trpm5*, a gene encoding an ion channel required for sweet, umami, and bitter taste signaling (Perez et al., 2002). Chromatin-accessible peaks were enriched in the promoter region and the transcription start site (TSS) only in type II taste cells. In contrast, the adjacent gene *Kcnq1* (Fig. 4b) showed high chromatin accessibility in its promoter and TSS regions in all three taste cell clusters (C1, C10, and C11 clusters), consistent with the expression of *Kcnq1* (encodes a pan-taste-cell marker) (Ohmoto et al., 2006; Wang et al., 2009a). In addition, co-accessibility analysis showed that many peaks in the surrounding regions of *Trpm5* were strongly co-accessible with the *Trpm5* peaks, suggesting these genomic regions may contain putative regulatory elements for *Trpm5* gene expression.

Similarly, the three *Tas1rs* that encode umami and sweet taste receptors showed accessible peaks selectively in type II taste cells (Fig. 4c). Furthermore, many other type II taste cell marker genes, such as *Gnat3*, *Plcb2*, *Pou2f3*, and *Calhm1*, also exhibited selective chromatin-accessible peaks in type II taste cells (Supplementary Fig. 4b).

Genes encoding markers for type I and III taste cells, such as *Entpd2*, *Kcnj1*, *Car4*, *Gad1*, and *Pkd2l1*, showed peaks selectively either in type I or type III taste cells (Supplementary Fig. 5). Interestingly, *Otop1*, which encodes a recently identified sour taste receptor (Teng et al., 2019), showed enriched accessibility in both type I and type III cell clusters (Fig. 5b). However, at least one peak within an intronic region of *Otop1* showed specific accessibility in type III cells. Again, such peaks could contain cis-regulatory elements for *Otop1* gene expression in type III taste cells.

**Fig. 5.**
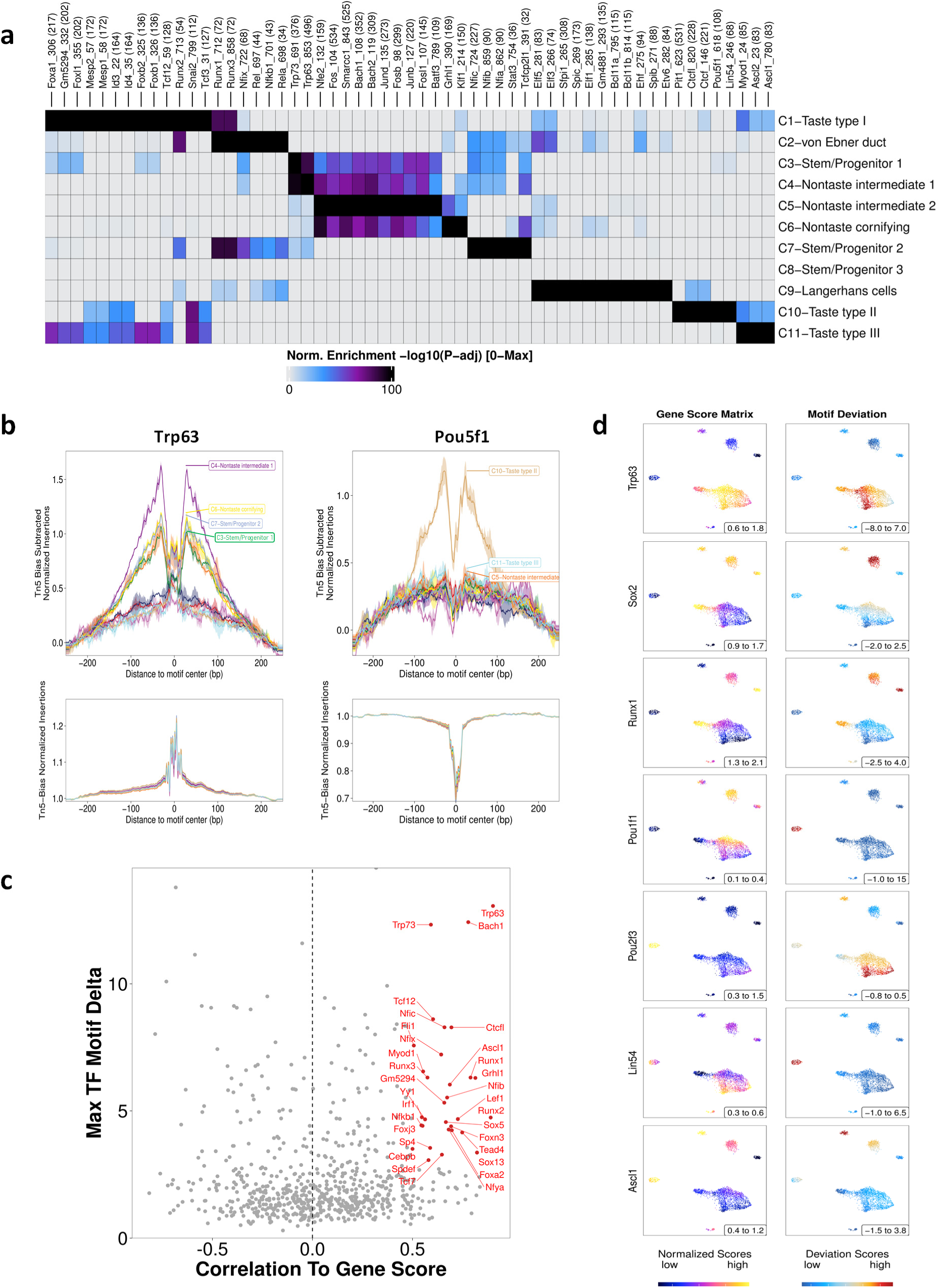
scATAC-seq revealed potential transcriptional regulators in taste bud cells. Transcription factor motif analyses in accessible peaks from control mice. **a.** A heatmap of enriched motifs of transcription factors across cell clusters. **b.** DNA footprints of Trp63 and Pou5f1 across cell clusters revealed by ArchR. **c.** Correlations between gene scores of transcription factors (TF) and their motif enrichment in cell clusters. Red indicates transcription factors with significant positive correlation, suggesting they likely act as transcription activators. **d.** Side-by-side comparisons of gene scores and motif enrichment of seven transcription factors. A lack of correlation between gene score and motif enrichment suggests the transcription factor may function as transcription repressor.

Together, our results revealed strong cell-type-selective chromatin accessibility for *Tas2rs* and genes of other receptor/signaling proteins involved in taste reception and signaling. The scATAC-seq data set can be used to identify putative cis-regulatory elements for taste-cell-specific gene expression, an understudied area of taste biology.

### Motif analyses identified potential transcriptional regulators in taste cells

An advantage of scATAC-seq is the ability to identify candidate transcriptional regulators. To do this, we performed binding motif enrichment analysis for transcription factors using accessible peaks (Supplementary Fig. 6a). This analysis identified many enriched motifs for an array of transcription factors (Fig. 5a and Supplementary Data 2). Notably, the binding motif for Trp63 was highly enriched in the C3 and C4 cell clusters (Fig. 5a and Supplementary Fig. 6f). This result is highly in line with previous reports that showed Trp63 is a marker for lingual epithelial stem/progenitor cells that give rise to both taste bud cells and nontaste epithelial cells (Bloomquist et al., 2019; Okubo et al., 2009), and that Trp63-deficient mice exhibit abnormal differentiation of the tongue epithelium (Mills et al., 1999). Trp63 footprint analysis using ArchR also showed enrichment in C3, C4, and C7 cell clusters (Fig. 5b).

Additionally, we identified several potential transcriptional regulators whose motifs were enriched in the C7 cluster of stem/progenitor cells, including Stat3, Tcfcp2l1 (i.e., Tfcp2l1), Nfia, and Nfib (Fig. 5a). All of these transcription factors are known to play critical roles in governing stemness or other qualities of stem cells (Chang et al., 2013; Raz et al., 1999; Tchieu et al., 2019).

Furthermore, we identified multiple potential regulators of taste bud cell differentiation and/or function (Fig. 5a). For instance, our results showed that the motifs of Ascl1, Ascl2, and Myod1 are enriched in type III taste bud cells (Fig. 5a and Supplementary Fig. 6f). Of these transcription factors, Ascl1 has been shown to play key roles in type III taste cell differentiation (Seta et al., 2011), while the functions of Ascl2 and Myod1 in type III taste cells remain to be determined. The motifs of Lin54, Pou5f1, Ctcf (CCCTC-binding factor), Ctcfl (Ctcf-like), and Pit1 (Pou1f1) were enriched in type II taste cells (Fig. 5a and Supplementary Fig. 6f). Of these transcription factors, Ctcf was shown to play a role in expression of human *TAS2R8* (Kojima et al., 2020). DNA footprints showed that Pou5f1, Lin54, and Pit1 are promising candidates as transcriptional regulators of type II taste cell (Fig. 5b and Supplementary Fig. 6b, c). Similarly, the motifs of multiple transcription factors are enriched in type I taste cells (Fig. 5a). Of these transcription factors, Foxa1 was reported as a potential target gene of Sonic hedgehog (Shh) (Golden et al., 2021), whose gene score was enriched in type I taste cells (Supplementary Data 1). The binding motif of Sox2 was also enriched in type I cells (Supplementary Fig. 6f and Supplementary Data 2). In addition, some of the identified transcription factors showed enriched motifs in all three taste cell clusters, such as Snai2, Tcf3, Id3, and Id4 (Fig. 5a and Supplementary Fig. 6f), suggesting they may play general roles in taste bud differentiation.

Transcription activators often show positive correlation between motif enrichment and gene expression, while transcription repressors often lack such correlation or show anti-correlation (Finkbeiner *et al*., 2022; Granja *et al*., 2021). Next, we performed correlation analysis of gene scores (proxies for gene expression) versus motif enrichment of transcription factors. Figure 5c shows the identified potential transcription activators. Trp63, Ascl1, and Nfib are among these factors, suggesting they may act as transcription activators in taste/lingual epithelium differentiation. Figure 5d and Supplementary Fig. 6g show side-by-side comparisons of gene scores and motif enrichments of various transcription factors. Notably, some of the transcription factors that have been shown to play important roles in taste cell differentiation, such as Pou2f3 and Nkx2.2, did not show correlation of gene score and motif enrichment, suggesting they may function as transcription repressors. This result is in agreement with a previous report that showed gene knockout of Pou2f3 resulted in expanded expression of type III cell-specific genes in type II taste cells (Matsumoto *et al*., 2011), suggesting Pou2f3 is a repressor of type III gene expression in type II taste cells (in addition to other possible functions). Together, our results revealed many potential transcriptional regulators in specific taste cell types.

### LPS induced broad epigenetic changes in taste tissue stem cells and taste bud cells

Next, we compared the scATAC-seq data sets from PBS- and LPS-treated mice. Figure 6a shows UMAP plots of the combined data set. The percentages of cells and accessible peaks in each cell cluster are shown in Fig. 6b and Supplementary Table 1. LPS treatment decreased the percentage of cells in the C7 cell cluster (11.36% in PBS vs. 8.09% in LPS data set) but increased the percentage of cells in the C8 cell cluster (0.18% in PBS vs. 4.44% in LPS data set). These results indicate that LPS strongly affects taste tissue stem/progenitor cells, consistent with our previous report (Cohn *et al*., 2010).

**Fig. 6.**
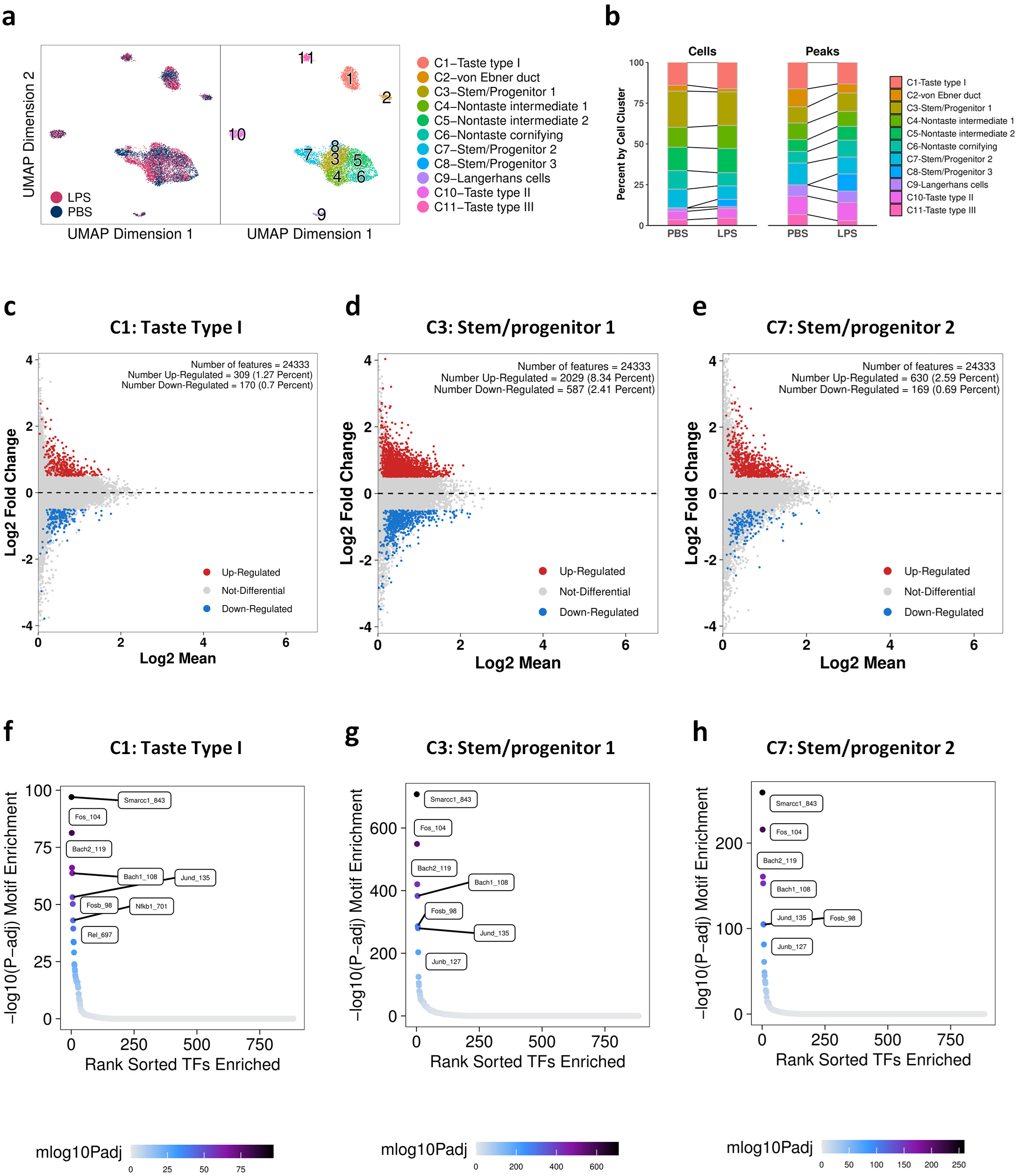
LPS induced broad epigenomic changes in taste bud cells and taste tissue stem/progenitor cells. **a.** UMAP plots. Left: Comparison of cells from control (PBS) and LPS-treated mice. Right: Cell clusters in the PBS and LPS combined data set. **b.** Percentages of cells and accessible peaks in each cell cluster. LPS treatment resulted in changes in the proportion of cells and peaks in some cell clusters. **c-e.** Up- and down-regulated genes (based on gene scores) in type I taste cells (**c**) and two clusters of taste tissue stem/progenitor cells (**d, e**). **f-h**. Up-regulated transcription factor (TF) motifs by LPS in type I taste cells (**f**) and two clusters of taste tissue stem/progenitor cells (**g, h**).

Gene score analysis showed many up- and down-regulated genes by LPS treatment. In particular, type I taste cells showed numerous up- and down-regulated genes (Fig. 6c). Pathway analysis of up-regulated genes revealed that many immune response pathways were significantly enriched, such as the TLR signaling pathways, cytokine production and signaling pathways, MAP kinase pathways, chemokine production, and immune cell migration pathways (Supplementary Fig. 7a). Consistently, motif analysis showed that LPS increased the accessible motifs for nuclear factor-κB (Nfkb1 and Rel; Fig. 6f), a master regulator of immune responses. These results strongly suggest that, in the presence of PAMPs, taste bud cells are capable of producing immune regulatory factors, such as cytokines and chemokines, consistent with our previous findings (Wang *et al*., 2007).

Furthermore, the C3 and C7 stem/progenitor cell clusters also showed strong inflammatory responses to LPS. We observed a large number of up- and down-regulated genes in these clusters (Fig. 6d, e). LPS significantly increased the accessible binding motifs for Smarcc1 (a subunit for the chromatin remodeling complex SWI/SNF), Bach1, Bach2, Fos, and JunD (subunits of AP-1 transcription factor known to be involved in LPS-induced gene expression) (Fig. 6g, h). These results suggest that LPS treatment induces large-scale chromatin remodeling in taste stem/progenitor cells. Pathway enrichment analysis showed up-regulation of defense pathways to LPS and bacteria, cytokine production and signaling, chemokine signaling and chemotaxis, MAP kinase cascade, and angiogenesis pathways – all related to inflammation (Supplementary Fig. 7b).

Interestingly, we observed some differences in pathway enrichment between the C3 and C7 clusters. TNF production, chemotaxis, and protein translation are more enriched in the C3 cluster, while interferon-γ production, IL-1 responses, antiviral pathways, and glucuronidation pathways are more enriched in the C7 cluster (Supplementary Fig. 7b). The significance of these differences remains to be determined.

### LPS increased chromatin accessibility at multiple *Tas2r* genomic loci

Next we performed detailed analysis of *Tas2r* genomic regions to evaluate whether LPS stimulation increased chromatin accessibility in these regions. As shown in Fig. 2d, expression of multiple *Tas2rs* in the genomic region between *Tas2r115* and *Tas2r125* on mouse chromosome 6 was induced >3-fold by LPS. Thus, we specifically compared the number of accessible peaks in this region between the PBS and LPS data sets. Figure 7a shows that the number of accessible peaks in this region in the LPS data set was more than double that in the PBS data set. Analysis of more mouse *Tas2rs* showed a general increase in the number of accessible peaks (Fig. 7b).

**Fig. 7.**
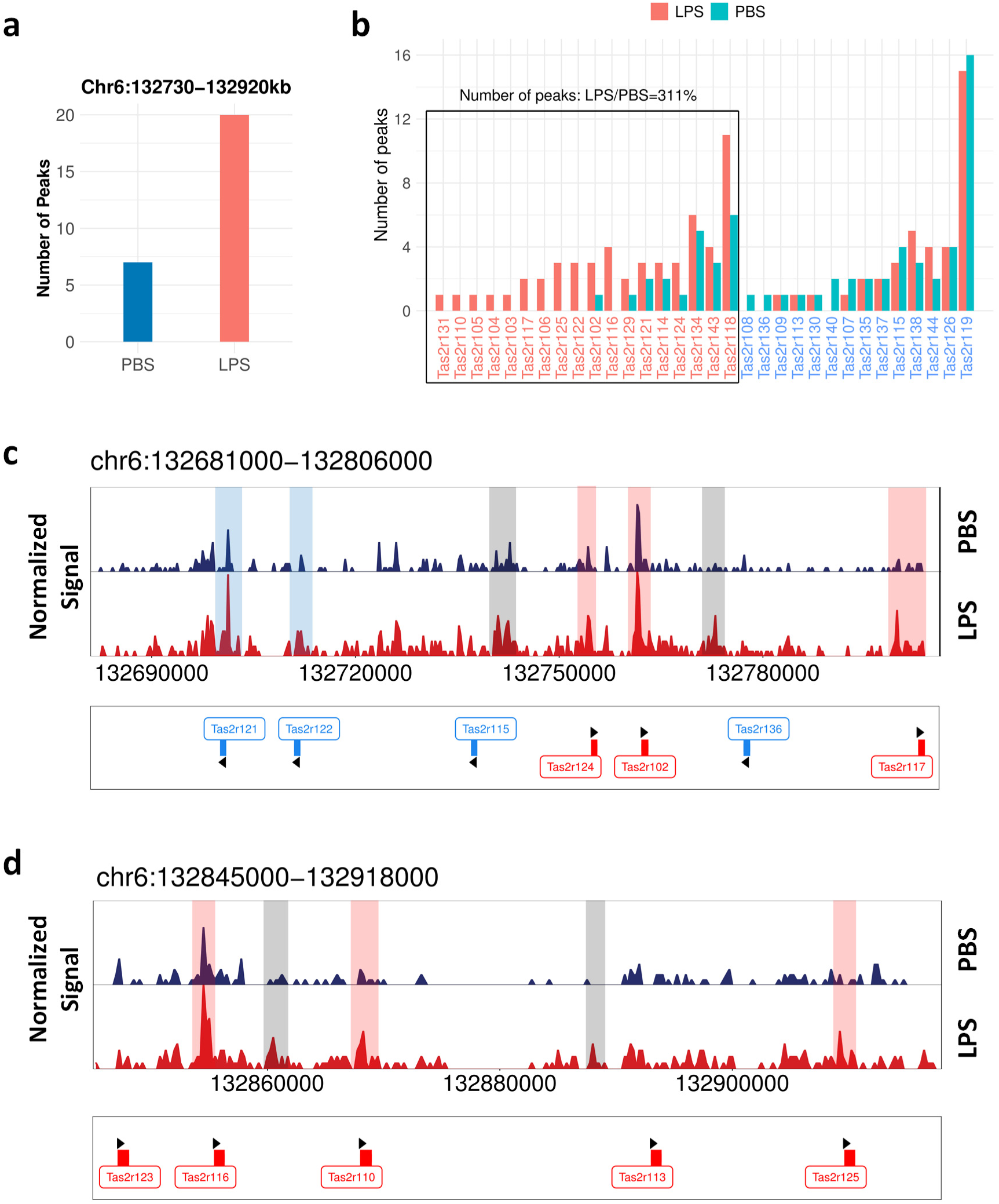
LPS increased chromatin accessibility of the majority of *Tas2rs*. Comparison of chromatin accessibility of the *Tas2r* loci between PBS- (control) and LPS-treated mice. **a.** Number of accessible peaks in the genomic region of the *Tas2r* gene cluster between *Tas2r115* and *Tas2r125* on chromosome 6 that harbors several LPS- highly inducible *Tas2rs*. LPS treatment markedly increased the number of peaks in this region. **b.** Number of accessible peaks for individual *Tas2rs*. LPS increased the number of peaks for the majority of *Tas2rs* (boxed area). **c, d.** Comparison of normalized peaks between the PBS and LPS groups in two *Tas2r* regions on chromosome 6. Gene symbols in red indicate genes transcribed in the forward direction, and their LPS-induced accessible peaks are highlighted in pink. Gene symbols in blue indicate genes transcribed in the reverse direction, and their LPS-induced accessible peaks are highlighted in blue. Gray bars highlight LPS-inducible peaks in intergenic regions. Black arrowheads indicate direction of *Tas2r* gene transcription.

We also compared normalized peak signals in the genomic region between *Tas2r121* and *Tas2r125* from PBS- and LPS-treated mice. In addition to the generally increased chromatin accessibility at the TSS of *Tas2rs*, increased accessibility was observed in some intergenic regions (Fig. 7c, d), suggesting these regions may contain cis-regulatory elements related to LPS-induced *Tas2r* gene expression. Together, these results show that LPS increases chromatin accessibility of the majority of the *Tas2rs*, indicating that epigenetic changes may underlie the increased *Tas2r* gene expression and elevated neural and behavioral responses to bitter compounds in this inflammation model.

## Discussion

In this study we showed that inflammation induced by LPS augmented bitter taste via epigenetic mechanisms that increased *Tas2r* gene expression. scATAC-seq identified many potential cis-regulatory regions in *Tas2rs* and other taste signaling genes and candidate transcriptional regulators in taste bud cells. We also observed extensive inflammatory responses in taste stem/progenitor cells and type I taste cells. Our study identified a previously unrecognized mechanism for modulating *Tas2r* gene expression and bitter taste.

Recent research has shown that taste GPCRs play important roles not only in the taste system but also in many extraoral tissues (Wang *et al*., 2021). T2Rs in solitary chemosensory cells are involved in detecting bacterial quorum-sensing molecules and triggering antimicrobial responses (Lee *et al*., 2014; Tizzano *et al*., 2010). LPS-induced *Tas2r* expression in taste buds could be part of the general immune response to a pathogen-derived molecule. Increased sensitivity to bitter compounds may enhance antimicrobial defense and alert the host to avoid exposure to toxins. The LPS-induced hyper-responsiveness to bitter compounds is consistent with our previous findings that showed knockout of TNF, a key inflammatory cytokine expressed by many immune cells as well as by taste bud cells, resulted in specific reduction in neural and behavioral responses to bitter compounds (Feng et al., 2015). Together, our studies provide strong evidence that bitter taste is selectively regulated by immune factors at the peripheral level.

Taste dysfunction contributes to loss of appetite. Elevated bitter taste could be a part of the anorexic responses associated with systemic inflammation. This mechanism may also contribute to rejection of bitter medicine, especially in young children. In this case, masking or down-regulating bitter taste could be beneficial. It is well established that LPS treatment reduces preference and intake of sweet taste compounds (Dantzer, 2001), which is considered a behavioral change related to depression caused by sickness (Dantzer et al., 2008).

Inflammatory cytokines, such as TNF and IL-1β, may play important roles in such behavioral changes by acting on various brain regions (Dantzer *et al*., 2008). Our behavioral and nerve recording results with sweet taste compounds (Fig. 1 and Supplementary Fig. 1) support these proposed brain mechanisms. Moreover, an intriguing recent study showed that fasting was protective in bacterial sepsis (Wang et al., 2016), suggesting anorexia may have important physiological roles during bacterial infections.

LPS-induced gene expression in macrophages can be separated into stages. Early primary response genes are induced quickly and do not require new protein synthesis, while secondary response genes are induced at a later time and largely depend on new protein synthesis and chromatin remodeling for induction (Medzhitov and Horng, 2009; Ramirez-Carrozzi et al., 2006). *Tas2rs* are likely secondary or even tertiary response genes due to their delayed induction (Fig. 2). In line with this, chromatin accessibility of the majority of *Tas2rs* was increased by LPS (Fig. 7), indicating remodeling of nucleosomes at *Tas2r* loci. Interestingly, COVID-19 infection also induces widespread changes in olfactory receptor gene expression (Zazhytska et al., 2022), providing another example of large-scale change in chemosensory receptor expression as an underlying cause for sensory dysfunction. Additionally, our data showed that some *Tas2rs*, which are close to one another on the chromosome, were co-accessible in type II cells and were likely co-regulated. We found several intergenic regions within the *Tas2r* gene clusters that showed selective accessible peaks in type II taste cells and were co-accessible with the surrounding *Tas2rs*. These intergenic regions may contain cis-regulatory elements, which will be determined in the future.

Emerging evidence shows that tissue stem cells play important regulatory roles in immune responses (Chen et al., 2019; Naik et al., 2018). Stem cells not only can produce cytokines and chemokines (Chen *et al*., 2019) but also can harbor epigenetic memory, which allows them to respond faster to future stimulation (Naik *et al*., 2018). Such immune memory is advantageous against future infections but may also contribute to chronic diseases. We observed that taste tissue stem/progenitor cells showed robust epigenetic responses to LPS treatment (Fig. 6), with up-regulation of numerous pathways related to cytokine/chemokine production and inflammation (Supplementary Fig. 7). The motifs of Smarcc1, a subunit of the chromatin remodeling complex SWI/SNF, were dramatically enriched in stem/progenitor cells after LPS induction, suggesting Smarcc1 may play key roles in LPS-induced chromatin remodeling in taste stem/progenitor cells. It is conceivable that chromatin remodeling may result in “inflammatory memory” in taste tissue stem/progenitor cells, leading to long-lasting effects on taste responses.

Surprisingly, von Ebner gland duct cells showed a robust immune regulatory signature, with high gene scores for an array of chemokines (Supplementary Fig. 3). Dclk1 is also enriched in these cells, suggesting this cell cluster may play important roles in immune surveillance, similar to tuft cells in the gut (Schneider et al., 2019). Salivary glands are major targets of the SARS-CoV-2 virus, which may, in part, contribute to taste loss associated with COVID-19 (Huang et al., 2021). Salivary gland duct cells may play a role in attracting immune cells to the proximity of taste tissues and contribute to taste tissue injury.

Our study identified many potential cell-type-selective transcriptional regulators (Fig. 5). Transcriptional regulation of taste receptors and signaling proteins remains poorly understood. Only a handful of transcription factors have been shown to play roles in taste cell differentiation. Consistent with previous publications, our results showed that the motifs and gene scores of Trp63 and Ascl1 were enriched in taste stem/progenitor cells and type III taste bud cells, respectively (Fig. 5d). Their binding motifs and gene scores are highly correlated (Fig. 5c), suggesting they are transcription activators. Conversely, Pou2f3, Pou5f1, and Pou1f1 did not show correlation between enrichment of gene scores and that of their binding motifs, suggesting they may act as transcription repressors. The phenotype of Pou2f3-knockout mice supports this proposition: lacking Pou2f3 resulted in expression of type III cell-specific genes in type II taste cells (Matsumoto *et al*., 2011). Further characterization of these transcription factors and identification of their downstream targets can help determine their roles in taste bud cell differentiation as well as in taste alterations associated with diseases.

### STAR Methods Mice

All procedures involving mice were performed according to protocols approved by the Monell Chemical Senses Center Institutional Animal Care and Use Committee. Adult male and female C57BL/6J mice were purchased from the Jackson Laboratory and housed in a climate-controlled environment at the animal facility of the Monell Chemical Senses Center. For induction of inflammation, LPS (purchased from Sigma, 2.5 mg/kg in PBS) or PBS (a vehicle control) was given by intraperitoneal injection.

### Brief-access behavioral tests

Brief-access tests were performed using Davis MS-160 mouse gustometers (Dilog Instruments, Tallahassee, FL) as previously described (Kim et al., 2012). Briefly, adult mice (age and sex matched) were trained for brief-access tests several days before LPS or PBS injection. The following taste compounds were tested: quinine hydrochloride (0.1, 0.3, and 3 mM) and sucrose (0.1, 0.2, and 0.6 M). To motivate sampling of taste solutions, mice were water-deprived for 22.5 h before the 30-min training sessions and the 30-min test sessions for quinine as a bitter taste compound and were food- and water-restricted (1 g of food and 1.5 ml of water) for 23.5 h before test sessions for sucrose as a sweet taste compound. In each test session, mice were tested with the three different concentrations of the taste compound along with a water control. Water and taste compounds were randomly presented to mice following random presentation schemes generated by the computer software. LPS-treated mice and control mice were tested at the same time in parallel: on day 2 and day 4 after LPS or PBS injection, mice were tested with sucrose solutions, and on day 3, with quinine solutions. For data analyses, lick ratios were calculated by dividing the number of licks for taste compounds by the number of licks for water presented in the same test session. For tests of quinine, aversion scores were calculated as 1 – lick ratio. We used repeated ANOVA to compare the between-group effect (PBS vs. LPS), with the taste stimulus concentrations treated as within-group effects, followed by post hoc t test.

### Gustatory nerve recordings

Chorda tympani and glossopharyngeal nerve recordings were carried out as described previously (Kim *et al*., 2012). Briefly, mice were anesthetized with pentobarbital (50-60 mg/kg of body weight, i.p.), and the trachea was cannulated. Mice were then fixed in the supine position with head holders. For recordings from the chorda tympani nerve, the right nerve was exposed at its exit from the lingual nerve and cut near its entrance to the bulla. For recordings from the glossopharyngeal nerve, the right nerve was exposed by removal of the digastric muscle and posterior horn of the hyoid bone and then dissected free from underlying tissues and cut near its entrance to the posterior lacerated foramen. For whole-nerve recording, the entire nerve was placed on a platinum wire recording electrode. An indifferent electrode was positioned nearby in the wound. Neural responses resulting from chemical stimulations of the tongue were fed into an amplifier (Grass Instruments, West Warwick, RI), monitored on an oscilloscope and an audio monitor. Whole-nerve responses were integrated with a time constant of 1.0 s and recorded using a computer for later analysis using a PowerLab system (PowerLab/sp4; AD Instruments, Colorado Springs, CO).

The following solutions were used as stimuli: 0.1-20 mM quinine hydrochloride, 0.1-30 mM denatonium, 0.003-0.5 mM cycloheximide, 3-300 mM MgSO_2_, 10-1000 mM sucrose, 30-1000 mM glucose, 0.3-30 mM sucralose, 0.3-20 mM saccharin, 0.1-10 mM inosine-5’-monophosphate (IMP), 10-1000 mM monopotassium glutamate (MPG) with or without 0.5 mM IMP, 10-1000 mM NaCl with or without 0.1 mM amiloride, 0.1-10 mM HCl, 1-100 mM citric acid, and 100 mM NH_4_Cl. These chemicals were dissolved in distilled water and used at ∼24°C. During recordings the test solutions were flowed for 30 s (chorda tympani recordings) or 60 s (glossopharyngeal recordings) at the same flow rate and temperature as distilled water used for rinsing the tongue (∼0.1 ml/s). The tongue was rinsed with distilled water for 1 min between successive stimulations.

To analyze whole-nerve responses to each stimulus, the magnitudes of integrated responses at different time points after stimulus onset were measured and averaged. The relative response magnitude for each test stimulus was calculated against the response magnitude to 100 mM NH_4_Cl, and this value was used for statistical analysis (ANOVA with post hoc t test) and for plotting dose-response curves. For chorda tympani recordings, 6-12 mice per group were used; for glossopharyngeal recordings, 5-10 mice per group were used.

### qRT-PCR

At various time points after PBS and LPS injection, mice were euthanized and tongue epithelium was peeled off as described previously (Wang *et al*., 2007). Circumvallate and foliate epithelia containing taste buds were cut and pooled together. Total RNA was extracted using Absolutely RNA Microprep Kit (Agilent) and then reverse transcribed into cDNA using Superscript III reverse transcriptase (Thermo Fisher Scientific). qPCR reactions were set up using Power SYBR Green PCR Master Mix (Thermo Fisher Scientific) in duplicate or triplicate and run on a StepOnePlus Real-Time PCR System (Thermo Fisher Scientific). Relative quantification of gene expression was performed using StepOnePlus software based on the 2^-ΔΔCt^ method. β-Actin was used as the endogenous control gene for these analyses. The specificity of the PCR reactions was analyzed by dissociation studies and confirmed by agarose gel electrophoresis. RT-PCR primers are listed in Supplementary Table 2. For statistical analysis, t test was conducted, with *p* > 0.05 considered significant.

### Isolation of nuclei, preparation of scATAC-seq libraries, and high-throughput sequencing

To isolate the nuclei for scATAC-seq, the peeled tongue epithelia from PBS- and LPS-treated mice were prepared as described previously(Wang *et al*., 2007). Six mice per group were used. Epithelia from the circumvallate papillae were cut off and rinsed in calcium-free Tyrode’s buffer (140 mM NaCl, 5 mM KCl, 10 mM HEPES, 1 mM MgCl_2_, 10 mM glucose, 10 mM Na-pyruvate, 2 mM EGTA, pH 7.4). Circumvallate epithelia were then transferred to a solution containing 0.25% trypsin-EDTA (Fisher Scientific) and incubated at 37°C for 15 min. Cold Opti-MEM with 10% fetal bovine serum (FBS) (Thermo Fisher Scientific) was added to the mixture, and cells were further dissociated by gentle pipetting. Cells were centrifuged, rinsed with cold Opti-MEM with 10% FBS, passed through a 40 µm cell strainer, and rinsed once with wash buffer (10 mM Tris-HCl, pH 7.5, 10 mM NaCl, 3 mM MgCl_2_, 1% bovine serum albumin (BSA), and 0.1% Tween-20). Dissociated cells were then resuspended in 0.1x Lysis buffer (10 mM Tris-HCl, pH 7.5, 10 mM NaCl, 3 mM MgCl_2_, 0.9% BSA, 0.01% Tween-20, 0.01% NP40, 0.001% digitonin) and incubated on ice for 5 min. Cell lysis was monitored by staining with trypan blue solution (0.4%; Thermo Fisher Scientific). Nuclei were rinsed with wash buffer, centrifuged, resuspended in 1x nuclei buffer (10x Genomics), and counted. About 4000 nuclei per group were used for downstream ATAC-seq procedures.

The chromatin transposition reaction, nuclei partitioning, barcoding, and library construction were carried out at the Next-Generation Sequencing Core (NGSC) facility at the University of Pennsylvania using 10x Genomics Chromium scATAC reagent kits and instruments, following the manufacturer’s protocols. High-throughput sequencing was also performed at the NGSC using Illumina HiSeq 4000 instruments.

### Data processing with Cell Ranger ATAC pipeline

Demultiplexed raw base call files generated from sequencing were used as inputs to the 10x Genomics Cell Ranger ATAC pipeline (version 1.2.0). FASTQ files were generated and aligned to the mm10 (i.e., GRCm38) mouse genome assembly using the Burrows-Wheeler Aligner Maximal Exact Matches (BWA-MEM) algorithm(Li and Durbin, 2010). Total numbers of read pairs were 236,312,328 bp and 246,253,784 bp for PBS- and LPS-treated mice, respectively. Fragment files were generated containing all unique properly paired and aligned fragments with mapping quality (MAPQ) > 30. The start and the end of the fragments were adjusted (4 bp from the leftmost alignment position and backward 5 bp from the rightmost alignment position) to account for the 9-bp overhang region that the transposase enzyme occupies during transposition. The adjusted positions represent the center point between these cuts, and this position is recorded as a cut site that represents a chromatin accessibility event.

Prior to further data analyses, we previewed the processed data with the Loupe Browser (10x Genomics), an interactive visualization software to show ATAC-seq peak profiles for scATAC-seq cell clusters, similar to the analyses described below.

### Data analyses with the ArchR pipeline

The downstream data analysis was conducted with ArchR (Granja *et al*., 2021) version 1.0.2 pipeline in R (version 4.1.3) using Ubuntu 20.04.4 LTS. Fragment files generated from the Cell Ranger ATAC pipeline were loaded into the ArchR pipeline to create arrow files. We filtered all scATAC-seq profiles to keep those that had at least 1000 unique nuclear fragments and a TSS enrichment score of 4. During creation of the arrow files, we generated the metadata and matrices, which included a “TileMatrix” (containing insertion counts across genome-wide 500-bp bins) and a “GeneScoreMatrix” that stores the predicted gene expression scores based on weighting of insertion counts in tiles close to a gene promoter (see below). Potential doublets (i.e., a single droplet containing more than one cell) were identified, which accounted for 3.5% and 4.2% of the cells from the PBS and LPS samples, respectively, and were filtered out.

To minimize batch effects, the ATAC-seq batches for the PBS and LPS samples were merged and calculated for the latent semantic index (LSI), which uses a term frequency (TF) that has been depth normalized to a constant (10,000) followed by further normalization with the inverse document frequency (IDF), and then the resultant matrix was log-transformed (i.e., log(TF – IDF)). Using the iterative LSI approach (Satpathy et al., 2019), we computed an initial LSI transformation on the most accessible tiles. We next applied Harmony (Korsunsky et al., 2019) to regress out the batch effect, resulting in a new harmonized co-embedding.

For clustering and visualization, we used a graph clustering approach (addCluster function) by including the dimensionality reduction matrix “Harmony” with other default parameters to identify clusters of the merged samples (PBS+LPS) and then visualized the scATAC-seq data using UMAP dimensionality reduction technique.

For gene score calculation, we used the addGeneScoreMatrix function and calculated a gene score matrix for each arrow file at the time of creation. Each sample was independently computed for the counts of each tile (genome-wide fragment bins) per cell. 500 bp of the tile size was used for binning counts prior to gene activity score calculation and the maximum counts per tile allowed is 4. The gene model used for weighting peaks for gene score calculation was "exp(-abs(x)/5000) + exp(-1)", where x is the stranded distance from the transcription start site of the gene, and exp and abs stand for exponential and absolute value, respectively. For sparse scATAC-seq data, the MAGIC imputation method (van Dijk et al., 2018) was used to impute gene scores by smoothing signal across nearby cells.

For cell cluster annotation, we used the gene scores from the merged data for the PBS and LPS samples as input matrix. Across all features, each cell cluster was compared to its own background group of cells (all other clusters) to determine if the given cell cluster retains significantly higher accessibility. A cutoff threshold of false discovery rate (FDR)-corrected *p* < 0.01 and log_2_-transformed fold change (log2FC) > 1.25 was applied to identify genes that appear to be uniquely active in each cell type (Supplementary Data 1). This gene list was used to manually annotate the cell types.

To calculate the Simpson diversity index, we employed Simpson’s index of diversity (*D*), which presents as (1 – *D*) and ranges between 0 and 1, with greater values indicating larger sample diversity. We used Simpson’s index to characterize the composition of all cells across the identified 11 cell types for the PBS and LPS samples separately.

For peak calling, we first created pseudo-bulk replicates to group single cells such that the data from each single cell were combined into a single pseudo-sample that resembled a bulk ATAC-seq experiment. We performed peak calling for each cluster independently to get high-quality, fixed-width, nonoverlapping peaks that represent the epigenetic diversity of PBS and LPS samples. Briefly, fragments from cells were grouped by broad cell class, and peaks were called on all cluster fragments using MACS2. Peaks from each cell type were then combined, merging overlapping peaks (overlapping peaks called within a single sample were handled using an iterative removal procedure) to form a union peak set, and a cell-by-peak matrix was constructed. This matrix was binarized for all downstream applications.

For co-accessibility analysis, we calculated correlations in accessibility between two peaks across many single cells to identify cell-type-specific peaks as being co-accessible, because these peaks are often all accessible together within a single cell type and often all not accessible in all other cell types. We applied the function addCoAccessibility to cells from the merged PBS and LPS data set and then to the PBS and LPS data sets separately, and reported data from the PBS data set only.

Differentially accessible peaks for each cell type between the PBS and LPS samples were determined by performing a two-sided Wilcoxon signed-rank test and selecting peaks that had log-transformed fold change > 0.5 and FDR-corrected *p* < 0.05.

For transcription factor motif enrichment analysis and footprinting, we used Catalog of Inferred Sequence Binding Preferences (CIS-BP) motifs obtained from chromVAR motifs mouse_pwms_v2 (Schep et al., 2017) to calculate motif positions using motifmatchr in ArchR. We first added the motif matrix in the object of the control PBS sample, next used differential testing to define the set of significantly differential peaks of all cell types for motif enrichment, and then performed hypergeometric enrichment of a given peak annotation within the defined marker peaks. We sorted out the motif for peaks with log-transformed fold change > 0.3 and FDR-corrected *p* < 0.5 as cell-type-specific motif enrichment. Footprinting plots of the cell-type-specific motifs were generated using a normalization method that subtracts the Tn5 bias from the footprinting signal, because the insertion sequence bias of the Tn5 transposase could lead to misclassification of transcription factor footprints.

To identify potential positive transcriptional regulators, we first identified deviant transcription factor motifs by performing a ChromVAR deviation analysis. The deviation score (Z-score) was predicted for enrichment of transcription factor activity on a per-cell basis from sparse chromatin accessibility data with the GC-content bias correction. Using the motif and gene score matrix, we identified transcription factors whose motif accessibilities were correlated with their own gene score. We filtered the positive regulators as those whose correlation between the motif enrichment and the gene score was >0.5 with an FDR-adjusted *p*-value < 0.01 and a maximum inter-cluster difference in deviation Z-score in the top quartile. This analysis was done with the PBS data set only.

To identify differentially regulated transcription factors by LPS, we statistically compared the control and LPS samples to identify the up- and down-regulators and motifs in differentially accessible peaks in the C1 (type I taste cells), C3 (taste stem/progenitor 1), and C7 (taste stem/progenitor 2) clusters. The cutoff thresholds log-transformed fold change > 0.5 and FDR-corrected *p* < 0.1 were applied in the two-sided Wilcoxon signed-rank tests.

### Pathway enrichment analysis

Pathway enrichment analyses were done as described previously (Reimand et al., 2019). Marker genes from each cell cluster (FDR < 0.01 and log2FC > 0.5) were analyzed separately. For analysis of LPS-induced changes in gene scores, up- and down-regulated genes (FDR < 0.05 and log2FC > 0.5 or < -0.5) were analyzed separately. Gene lists were first analyzed in g:Profiler, excluding electronic Gene Ontology (GO) annotations and limiting gene terms to 5-1000. Enriched pathways identified by g:Profiler from GO molecular functions and GO biological processes were imported into Cytoscape software (version 3.9.1) and visualized using EnrichmentMap. Functionally connected pathways were further manually organized into groups for better visualization and interpretation.

## Data availability

Sequencing data generated in this study are available in GEO with accession no. GSExxxxx.

## Code availability

All custom R code is available at https://github.com/Cailu086Lin/scATACseq_mouse_taste.

## Acknowledgements

This study was supported in part by National Institutes of Health/National Institute of Deafness and Other Communication Disorders (NIH/NIDCD) grants R01DC018042 (H.W.), R42DC017693 (D.R.R.), and P30 DC011735 (R.F.M.). We also acknowledge funding from NIH G20OD020296 (D.R.R.; an infrastructure improvement grant at the Monell Chemical Senses Center).

## Author Contributions

C.L., M.J., J.Q., S.F., M.Z., and H.W. performed the experiments and analyzed the results. L.H. provided PCR primers for *Tas2rs*. Y.N., R.F.M., D.R.R., and H.W. co-supervised nerve recording experiments and/or scATAC-seq data analyses. All authors were involved in data discussion and results interpretation. C.L., P.J., I.M., L.H., R.F.M., D.R.R., and H.W. were involved in drafting or editing the manuscript.

---

**Supplementary Figure 1.**
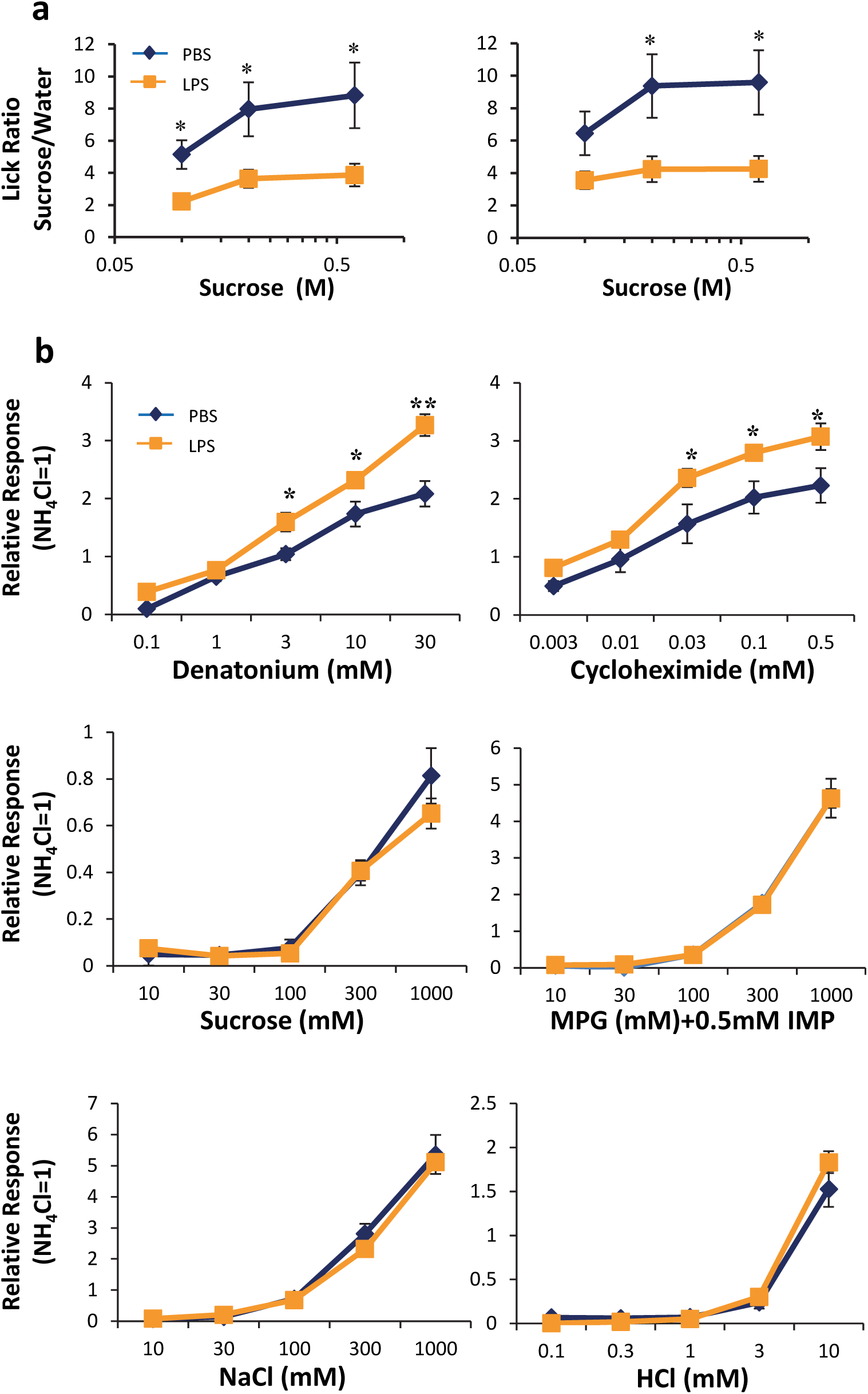
LPS induced neural and behavioral changes. **a.** LPS decreased preferences to sucrose in brief-access tests conducted on day 2 (left) and day 4 (right) after LPS or PBS injection. **b.** Glossopharyngeal nerve responses. LPS selectively increased responses to bitter taste compounds. MPG, monopotassium glutamate; IMP, inosine-5’-monophosphate. Data are means ± SE. N=5-10 mice per group.

**Supplementary Figure 2.**
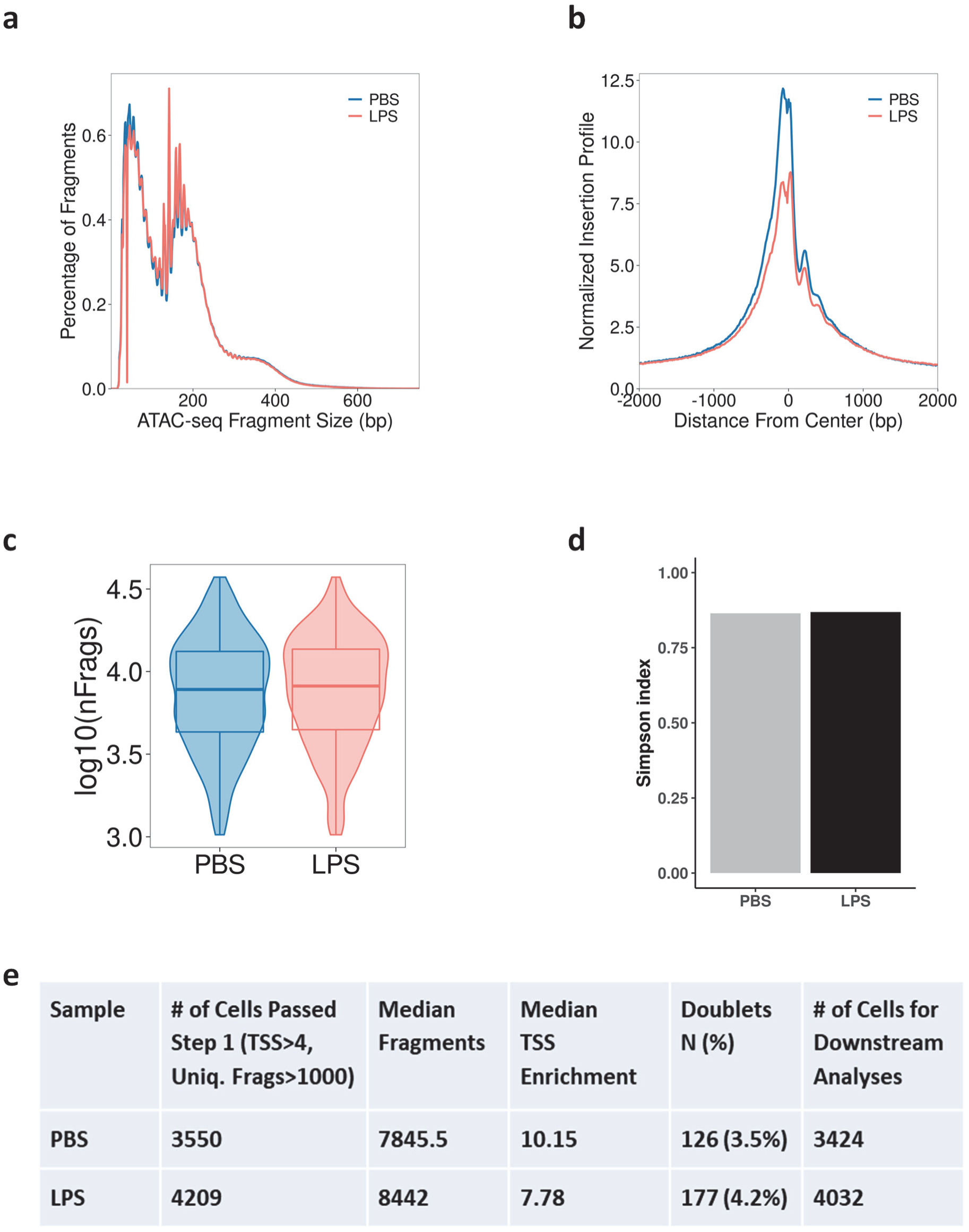
Quality check of the scATAC-seq data sets. **a.** scATAC-seq fragment size distribution, showing enriched DNA fragments for single or double nucleosomes. **b.** Enrichment of transposition insertion sites around transcription start sites. **c.** Distribution of the number of fragments per cell. **d.** Simpson diversity index. **e.** Summary table of quality check data. TSS, transcription start site.

**Supplementary Figure 3.**
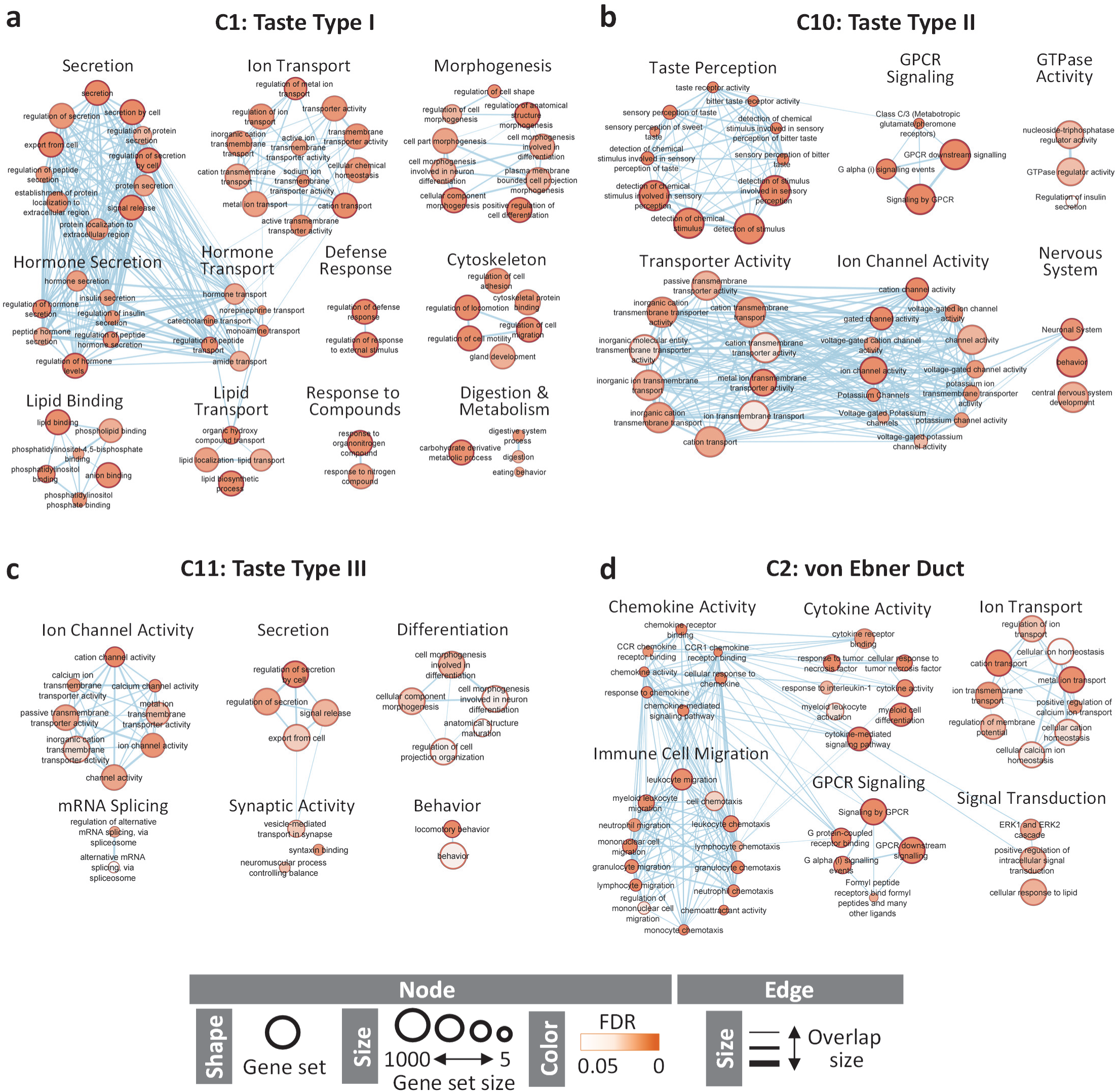
scATAC-seq revealed enriched biological pathways in taste bud cells and von Ebner gland duct cells. Identified marker genes from the indicated cell clusters from control mice were analyzed using g:Profiler and Cytoscape/EnrichmentMap software. **a.** An enrichment map of type I taste bud cells. Protein secretion, hormone secretion, and hormone transport are among the enriched pathways. **b.** An enrichment map of type II taste bud cells. GPCR signaling and taste perception are among the enriched pathways. **c.** An enrichment map of type III taste bud cells. Ion channel and synaptic activities are among the enriched pathways. **d.** An enrichment map of von Ebner gland duct cells showing an enriched immune regulatory signature.

**Supplementary Figure 4.**
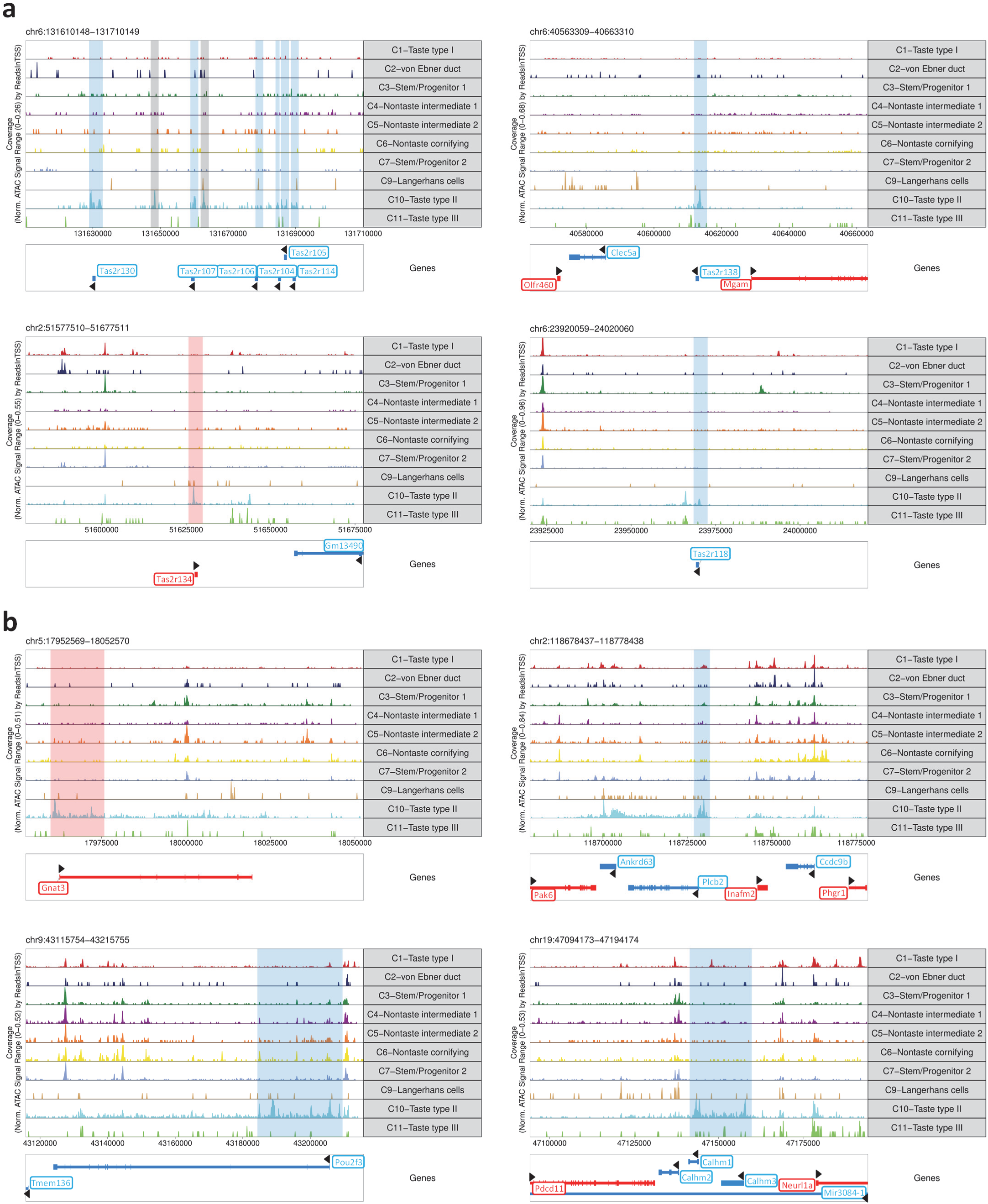
Chromatin-accessible peaks in four *Tas2r* genomic regions (a) and in the genomic regions of *Gnat3*, *Plcb2*, *Pou2f3*, and *Calhm1* (b). scATAC-seq data set from control mice. Gene symbols in red indicate genes transcribed in the forward direction, and their cluster-selective accessible peaks are highlighted in pink. Gene symbols in blue indicate genes transcribed in the reverse direction, and their cluster-selective accessible peaks are highlighted in blue. Gray bars in **a** indicate type II cell-selective peaks in intergenic regions. Black arrowheads also indicate direction of gene transcription.

**Supplementary Figure 5.**
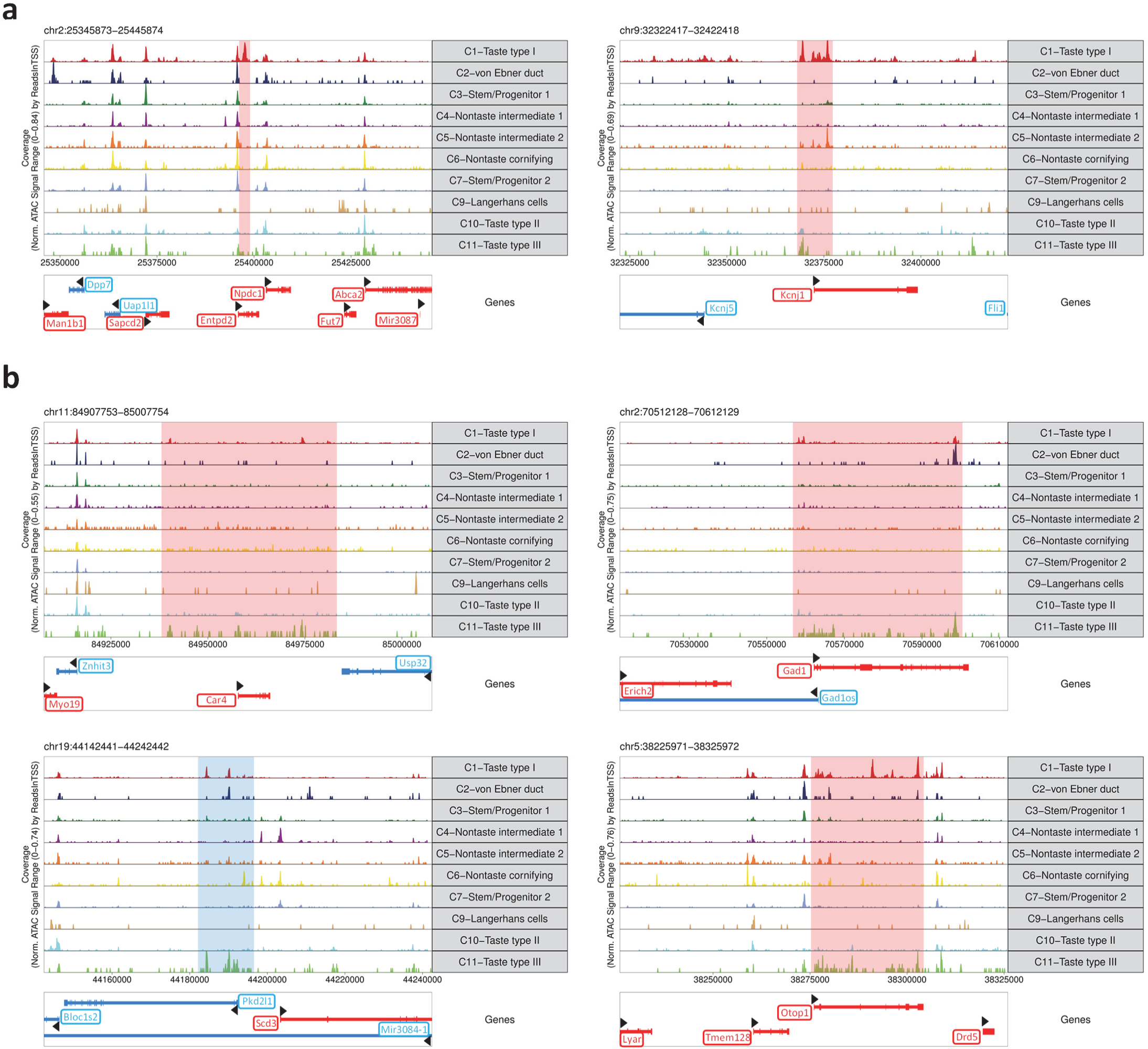
Chromatin-accessible peaks in the genomic regions of type I taste cell markers Entpd2 and Kcnj1 (a) and type III taste cell markers Car4, Gad1, Pkd2l1, and Otop1 (b). scATAC-seq data set from control mice. Gene symbols in red indicate genes transcribed in the forward direction, and their cluster-selective accessible peaks are highlighted in pink. Gene symbols in blue indicate genes transcribed in the reverse direction, and their cluster-selective accessible peaks are highlighted in blue. Black arrowheads also indicate direction of gene transcription.

**Supplementary Figure 6.**
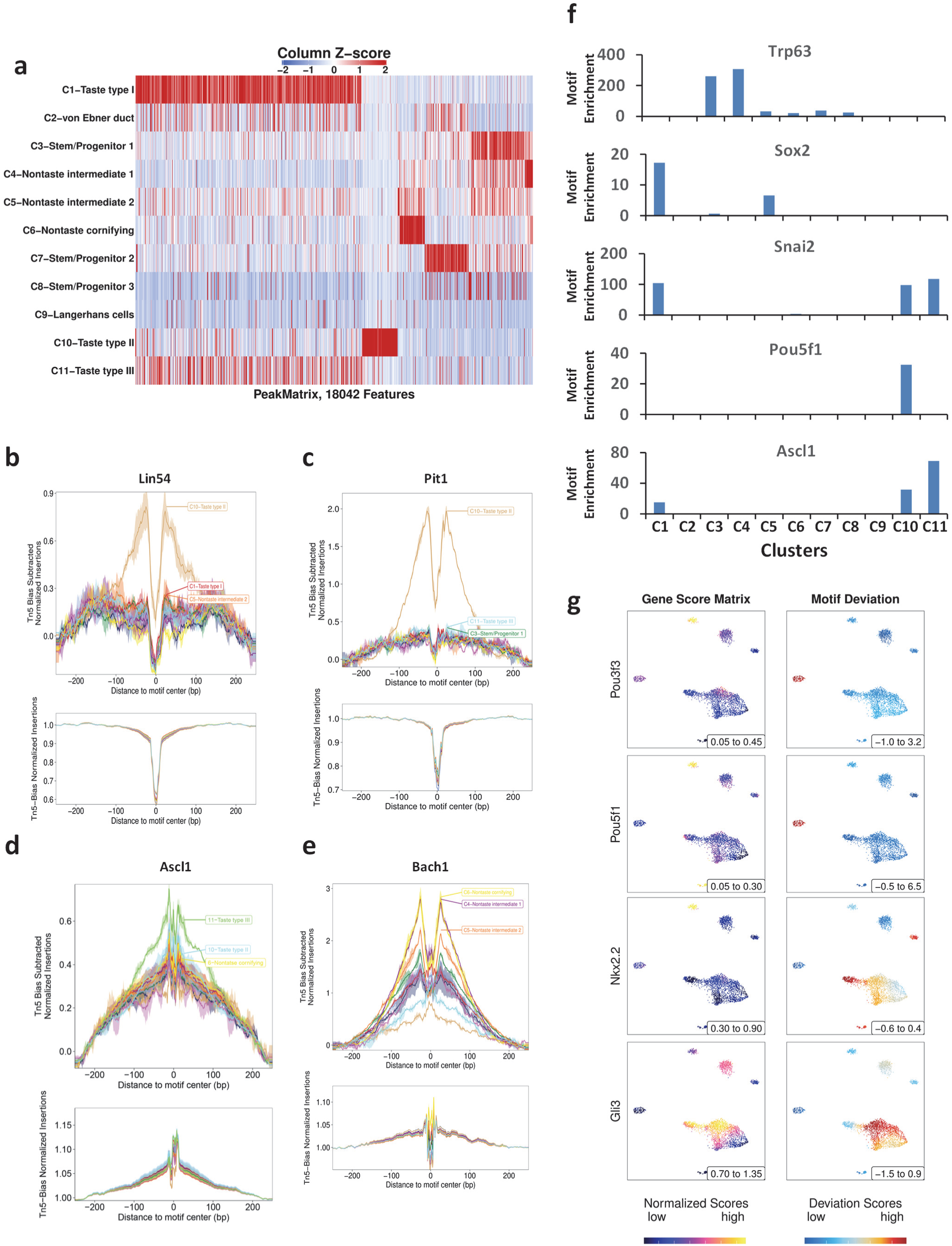
Potential transcriptional regulators. scATAC-seq data set from control mice. **a.** A heatmap of chromatin-accessible peaks by clusters. **b-e.** DNA footprints of Lin54, Pit1, Ascl1, and Bach1. **f.** Motif enrichment scores of Trp63, Sox2, Snai2, Pou5f1, and Ascl1 by cell cluster. **g.** Side-by-side comparisons of gene scores and motif enrichment of Pou3f3, Pou5f1, Nkx2.2, and Gli3.

**Supplementary Figure 7.**
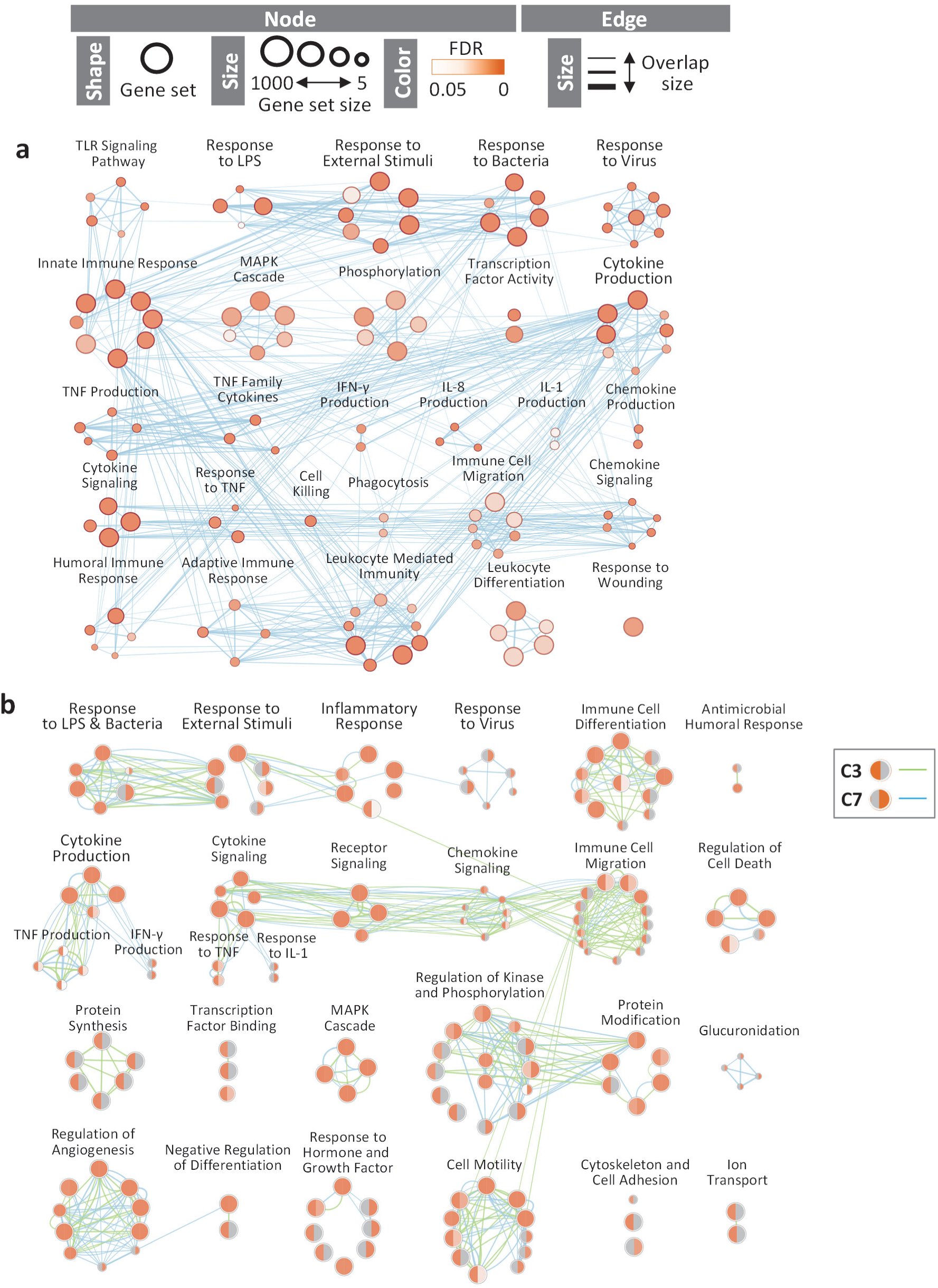
LPS induced inflammatory pathways in type I taste bud cells and taste tissue stem/progenitor cells. Genes up-regulated by LPS in type I taste cells (**a**) and two clusters of taste tissue stem/progenitor cells (**b**) were analyzed for pathway enrichment using g:Profiler and Cytoscape/ EnrichmentMap software. Each node (circle) represents a distinct pathway. Similar pathways are grouped and labeled according to their shared functions (pathway labels for individual nodes are not shown). The size of the node indicates the number of genes in the GO pathway. The color of the node represents the FDR value. Edges (blue or green) represent the number of genes overlapping between two pathways; thicker edges indicate more overlapping genes. In **b**, the C3 and C7 taste tissue stem/progenitor clusters are shown together, with the left halves of the nodes representing C3 and the right halves representing C7. Edges for C3 and C7 are in green and blue, respectively.

**Supplementary Table 1.** **The number and percentage of cells in each cell clusters in PBS and LPS scATAC-seq data sets.**

| Clusters | Annotation | # of Cells | % By Cluster | Sample | # of Cells | % By Sample |
| --- | --- | --- | --- | --- | --- | --- |
| C1 | Taste type I | 1128 | 15.13 | PBS | 479 | 13.99 |
|  |  |  |  | LPS | 649 | 16.10 |
| C2 | von Ebner duct | 205 | 2.75 | PBS | 125 | 3.65 |
|  |  |  |  | LPS | 80 | 1.98 |
| C3 | Stem/Progenitor 1 | 1590 | 21.33 | PBS | 759 | 22.17 |
|  |  |  |  | LPS | 831 | 20.61 |
| C4 | Nontaste intermediate 1 | 987 | 13.24 | PBS | 418 | 12.21 |
|  |  |  |  | LPS | 569 | 14.11 |
| C5 | Nontaste intermediate 1 | 1090 | 14.62 | PBS | 491 | 14.34 |
|  |  |  |  | LPS | 599 | 14.86 |
| C6 | Nontaste cornifying | 715 | 9.59 | PBS | 389 | 11.36 |
|  |  |  |  | LPS | 326 | 8.09 |
| C7 | Stem/Progenitor 2 | 725 | 9.72 | PBS | 392 | 11.45 |
|  |  |  |  | LPS | 333 | 8.26 |
| C8 | Stem/Progenitor 3 | 185 | 2.48 | PBS | 6 | 0.18 |
|  |  |  |  | LPS | 179 | 4.44 |
| C9 | Langerhans cells | 117 | 1.57 | PBS | 67 | 1.96 |
|  |  |  |  | LPS | 50 | 1.24 |
| C10 | Taste type II | 413 | 5.54 | PBS | 178 | 5.20 |
|  |  |  |  | LPS | 235 | 5.83 |
| C11 | Taste type III | 301 | 4.04 | PBS | 120 | 3.50 |
|  |  |  |  | LPS | 181 | 4.49 |

**Supplementary Table 2.**
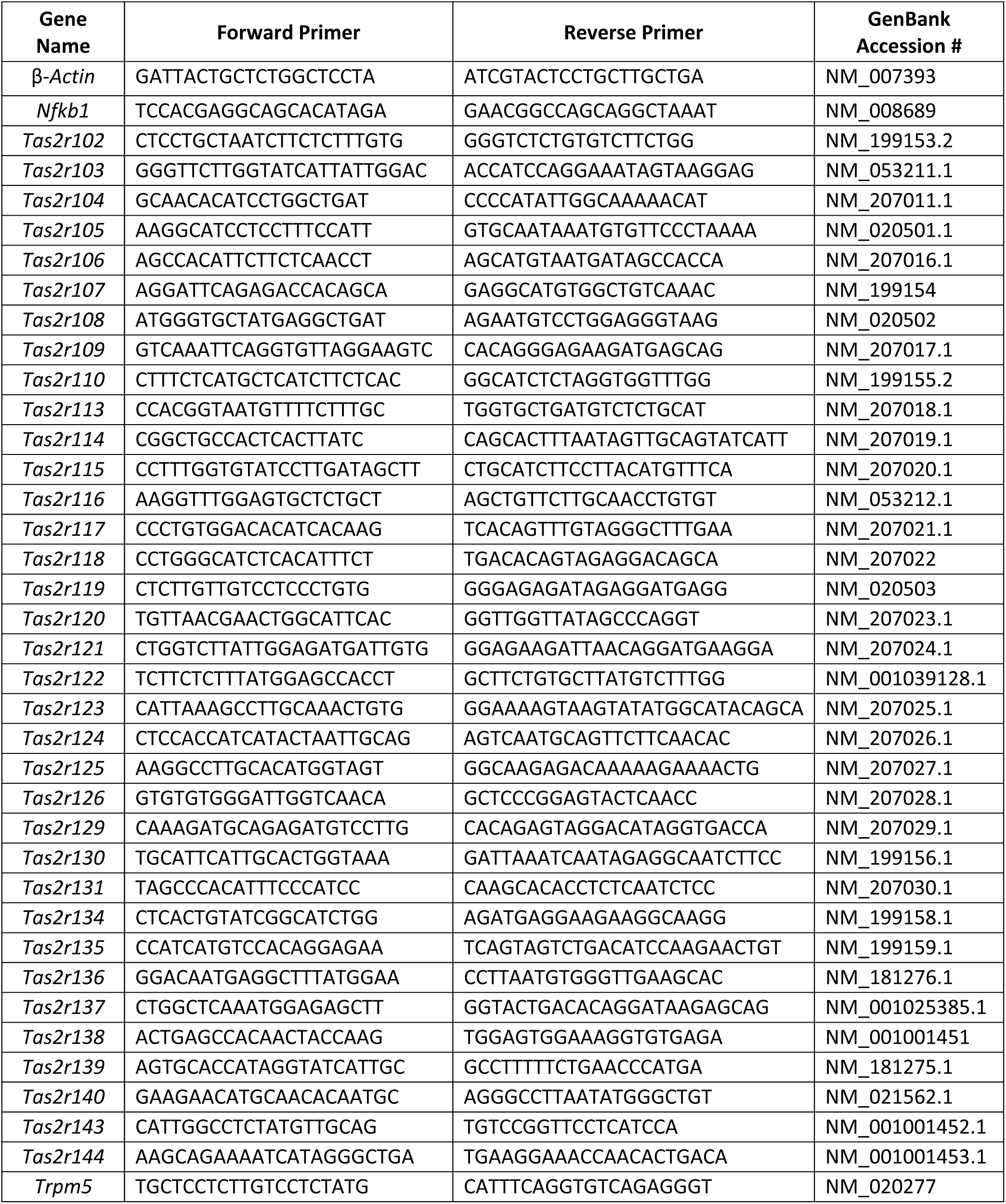
**Real-time PCR primers**

## Supplementary Data in Excel files

**Supplementary Data 1.** Gene scores of cell cluster marker genes from the control data set.

**Supplementary Data 2.** Enriched motifs by clusters from the control data set.

